# Deconstructing glucose-mediated catabolite repression of the *lac* operon of *Escherichia coli*: II. Positive feedback exists and drives the repression

**DOI:** 10.1101/2020.06.23.166959

**Authors:** Ritesh K. Aggarwal, Atul Narang

## Abstract

The expression of the *lac* operon of *E. coli* is subject to positive feedback during growth in the presence of gratuitous inducers, but its existence in the presence of lactose remains controversial. The key question in this debate is: Do the lactose enzymes, Lac permease and β-galactosidase, promote accumulation of allolactose? If so, positive feedback exists since allolactose does stimulate synthesis of the lactose enzymes. Here, we addressed the above question by developing methods for determining the intracellular allolactose concentration as well as the kinetics of enzyme induction and dilution. We show that during *lac* induction in the presence of lactose, the intracellular allolactose concentration increases with the lactose enzyme level, which implies that lactose enzymes promote allolactose accumulation, and positive feedback exists. We also show that during *lac* repression in the presence of lactose + glucose, the intracellular allolactose concentration decreases with the lactose enzyme levels, which suggests that under these conditions, the positive feedback loop turns in the reverse direction. The induction and dilution rates derived from the transient data show that the positive feedback loop is reversed due to a radical shift of the steady state induction level. This is formally identical to the mechanism driving catabolite repression in the presence of TMG + glucose.

## 1. Introduction

Two small molecules, namely 3’,5’-cyclic adenosine monophosphate (cAMP) (1) and allolactose (2) are known to regulate the expression of the *lac* operon of *Escherichia coli*. Mechanisms involving these molecules, namely cAMP-mediated transcriptional inhibition (3) and EIIA^glc^-mediated inducer exclusion (4), are thought to cause glucose-mediated *catabolite repression* (5–7), a 600-fold repression of the *lac* operon observed in the presence of glucose (8).

There is growing evidence that cAMP-mediated regulation, by itself, cannot account for the 600-fold repression (9). Indeed, it was widely believed earlier that this repression occurred due to the pronounced decline of cAMP levels in the presence of glucose (10). However, subsequent experiments showed that the intracellular cAMP levels during exponential growth on glucose and lactose were essentially the same, addition of exogenous cAMP to the medium did little to relieve the repression, and diauxic growth persisted in cAMP-independent mutants (8). Finally, the variation of steady state *lac* expression attributed to cAMP regulation (10) is due to enzyme dilution due to growth, and cAMP regulation *per se* changes the *lac* expression rate no more than 3-fold (11).

The evidence against the cAMP-mediated mechanism led to the hypothesis that inducer exclusion is the main mechanism of catabolite repression (8, 12, 13). This hypothesis was based on experiments showing that inducer exclusion is necessary for catabolite repression since abolishing inducer exclusion eliminated catabolite repression. Specifically, no catabolite repression is observed if a medium containing glucose and lactose is inoculated with wild-type cells plus high concentrations of IPTG, or mutant cells lacking functional repressor (LacI) or overexpressing the permease (LacY). However, other experiments show that inducer exclusion is not sufficient for catabolite repression since it decreases the permease activity only 2-fold in *lac*-constitutive cells (14), and no more than 6-fold in inducible cells (15). Thus, neither cAMP-mediated regulation nor inducer exclusion, by themselves, account for the 600-fold glucose-mediated *lac* repression.

We have proposed that glucose-mediated *lac* repression occurs because induction is subject to positive feedback which amplifies the small effects of cAMP-mediated regulation and inducer exclusion (16–19). Positive feedback occurs because lactose permease (LacY) and β-galactosidase (LacZ), referred to hereafter as *lactose enzymes*, promote the accumulation of allolactose, which in turn stimulates the synthesis of even more lactose enzymes. In the presence of lactose, this positive feedback loop results in the progressive accumulation of allolactose and the lactose enzymes. However, upon addition of glucose to a culture growing on lactose, the positive feedback loop turns in the *reverse* direction, thus leading to the progressive *depletion* of allolactose and the lactose enzymes. More precisely, upon the addition of glucose, the *lac* expression rate decreases due to the rapid decline of the cAMP and allolactose levels. The resultant decrease of the lactose enzyme levels causes the decline of the allolactose level, which in turn leads to further reduction of the lactose enzyme levels. The repetition of this “reversed cycle” results in the progressive depletion of both lactose enzyme and allolactose levels, ultimately yielding almost complete repression. Thus, *lac* induction and repression result from the turning of the same positive feedback loop, but in opposite directions.

Our model is formally identical to the models proposed for *lac* induction (20–22) and glucose-mediated repression in the presence of the non-metabolizable inducer TMG (15). However, theoretical studies have argued that the positive feedback loop postulated by our model exists only in the presence of non-metabolizable inducers — it does not exist in the presence of the metabolizable substrate lactose (23–25). Specifically, these studies acknowledge that both TMG and allolactose stimulate synthesis of the lactose enzymes, but these enzymes promote the accumulation of only TMG, and not allolactose. The mathematical basis of this claim is detailed in supplemental section S1, but it can be explained intuitively by considering the effect of increasing lactose enzyme levels on the intracellular TMG and allolactose concentrations, which can be assumed to be in quasi-steady state since they evolve much faster than the lactose enzyme levels. First, consider the case of cells exposed to TMG, which is accumulated in the cell by the permease, and depleted by passive diffusion (Fig. S1). If the permease level of such cells is increased, the accumulation rate of intracellular TMG is enhanced. Since intracellular TMG reaches quasi-steady state within minutes, this increase of the TMG accumulation rate must be rapidly followed by an equal increase of its depletion rate by passive diffusion. This can be achieved only if the intracellular TMG level increases by an amount proportional to the original increment of the permease level. Thus, the quasi-steady state intracellular TMG level is proportional to the permease level, which is a precise expression of the informal claim that the lactose enzymes promote the accumulation of TMG. Next, consider the case of cells exposed to intracellular lactose, which is accumulated by the permease, and depleted by β-galactosidase2 (Fig. S2). If the permease level of such cells is increased, the lactose uptake rate increases. However, since permease and β-galactosidase are co-ordinately synthesized, any increase of the permease level must be accompanied by a proportional increase of the β-galactosidase level. Consequently, the increase of the lactose uptake rate is automatically balanced by an equal increase of the lactose removal rate, and quasi-steady state is achieved without any change in the intracellular lactose level. The quasi-steady state intracellular lactose concentration is therefore independent of the lactose enzyme levels. Since allolactose is synthesized and depleted by the very same enzyme β-galactosidase (Fig. S2), it follows *a fortiori* that the quasi-steady state allolactose level is also independent of the lactose enzyme level. Thus, the lactose enzymes fail to promote *any* accumulation of allolactose, and hence there is no positive feedback at all.

The purported absence of positive feedback in the presence of lactose is based on two model assumptions that are inconsistent with the data and relaxing them restores the positive feedback. First, the models neglect the significant excretion of intracellular lactose and allolactose (26), which can restore the strong dependence of the quasi-steady state allolactose levels on the prevailing lactose enzyme levels. To see this, observe that the drastically different behaviour in the presence of TMG and lactose stems from model assumptions for TMG and lactose that represent two ends of a continuous spectrum: TMG, which is assumed to excreted but not metabolized, yields positive feedback, whereas lactose, which is assumed to be metabolized but not excreted, yields no positive feedback (27). If lactose is not only metabolised, but also excreted, it is conceivable that the behaviour in the presence of lactose approximates that in the presence of TMG (28). This intuitive argument is corroborated by mathematical analysis (Section S1), which shows furthermore that excretion of allolactose can also generate positive feedback. Second, the models assume that the LacY:LacZ ratio is always constant due to coordinate synthesis, but this is true only in the absence of inducer exclusion. In the presence of inducer exclusion, some of the LacY, but not LacZ, molecules are inactivated, so that the LacY:LacZ ratio is not constant, but increases with the LacZ level (15). Under this condition, the intracellular lactose and allolactose levels are not independent of, but increase with, the LacZ level, and positive feedback can exist in the presence of lactose (Section S1). As a side note, we may also observe that since LacY, but not LacZ, is partially inactivated in the presence of glucose, the activity of LacZ, rather than LacY, provides a more accurate measure of *lac* expression. Henceforth, the terms *induction level* and *induction rate* will refer to the specific activity and synthesis rate of LacZ, respectively.

The above argument shows that in theory, positive feedback can exist during growth on lactose, and hence, drive glucose-mediated repression, but there is no empirical evidence for this. Thus, the goal of this work is to experimentally test the validity of our model hypotheses:

1. Positive feedback exists, and drives glucose-mediated catabolite repression.
2. Positive feedback exists because allolactose stimulates lactose enzyme synthesis, and lactose enzymes promote allolactose accumulation.

To this end, we developed methods for determining the intracellular allolactose concentration as well as the induction and dilution rates of β-galactosidase during the course of *lac* induction in the presence of lactose and *lac* repression in the presence of lactose + glucose. We verified the first hypothesis by showing that positive feedback exists because the induction rate increases with the induction level, and positive feedback drives the repression because the induction and dilution rates in the presence of glucose + lactose are formally similar to those observed during catabolite repression in the presence of glucose + TMG (15). Insofar as our second hypothesis is concerned, it is clear that since allolactose is known to stimulate lactose enzyme synthesis (2), it suffices to check if the concentration of intracellular allolactose increases with the induction level. Two methods are generally used to quantify the intracellular concentrations of metabolites, namely the *cell-separation method* in which the concentration of the intracellular metabolite is measured after separating the cells from the medium, and the *difference method* in which the intracellular metabolite concentration is determined (without separating the cells from the medium) by subtracting the amount of metabolite in the medium from the total amount in the culture sample (containing both cells and medium) (29–33). However, both methods are prone to large errors when significant quantities of the metabolite accumulate in the medium (34, 35). We confirmed that there was significant accumulation of allolactose in the medium, and the above methods led to unacceptably large errors in the measurement of the intracellular allolactose concentration. Now, it turns out that the intracellular concentrations of several excreted compounds, such as cAMP and synthetic galactosides, are proportional to their specific efflux rates (10, 14, 36, 37). We therefore devised a new method, referred to hereafter as the *indirect* method to distinguish it from the *direct* methods described above, which exploits the allolactose efflux rate as a measure of its intracellular concentration. With the help of this indirect method, we measured the evolution of the intracellular allolactose levels during the course of induction in the presence of lactose, and repression in the presence of lactose + glucose. In both cases, the intracellular allolactose level increased with the induction level, which contradicts the conclusion derived from mathematical models, and verifies our second hypothesis. In the first case, the intracellular allolactose and induction levels increased in tandem, and in the second case, both levels decreased in tandem, which confirms that the turning of the positive feedback loop is reversed in catabolite repression. We show that this reversal is precipitated by a dramatic shift of the steady state induction level in the presence of glucose.

## 2. Material and Methods

### 2.1 Growth Conditions

The wild-type strain *E. coli* K12 MG1655 was obtained from Coli Genetic Stock Center (CGSC) at Yale University. Single colonies stored on LB Agar plates were inoculated in LB growth medium, grown for 6–8 hours at 37 °C, and then grown overnight at 37 °C after a 1:1000 dilution to M9 medium (38) supplemented with the appropriate carbon source. In the induction experiment, cells were pre-grown in maltose (5.6 mM), and transferred to pre-warmed medium containing maltose (2.8 mM) + lactose (4 mM). We used this mixture instead of pure lactose to minimize the heterogeneity of the cell population that occurs during growth on pure lactose (39). In the repression experiment, cells were pre-grown on maltose (2.8 mM) + lactose (4 mM), and transferred to pre-warmed medium containing glucose (2.2 mM) + lactose (4 mM). The cell density (gdw/l) was followed by measuring the optical density at 600 nm (OD_600_), which provided the cell density via the expression gdw/l = 0.35×OD_600_. To ensure saturating substrate levels throughout the experiment, partial removal of cells was often necessary. To this end, a part of the culture was centrifuged and the supernatant was reintroduced into the culture medium.

### 2.2 Sample preparation

For each time point, three aliquots of 0.5–2 ml each were withdrawn from the experimental cultures. The first aliquot was used to measure the optical density at 600 nm. The second aliquot, which was used to measure the β-galactosidase activity, was stored in an ice-water bath until the assay. The third aliquot, which was used to measure extracellular cAMP and allolactose concentrations, was rapidly filtered through a 0.22 μm nylon or polyethersulfone filter, and transferred to a boiling water bath to deactivate any residual enzymes. These filtered samples (also referred to as *medium* samples) were appropriately diluted in deionized water and stored at −20 °C until cAMP and/or sugar analysis. For measuring the total allolactose concentrations (referred to as *total* samples in Fig. 1), a fourth aliquot was also drawn, which was rapidly diluted in boiling water to deactivate enzymes and release the intracellular allolactose into the medium.

**Fig. 1.**
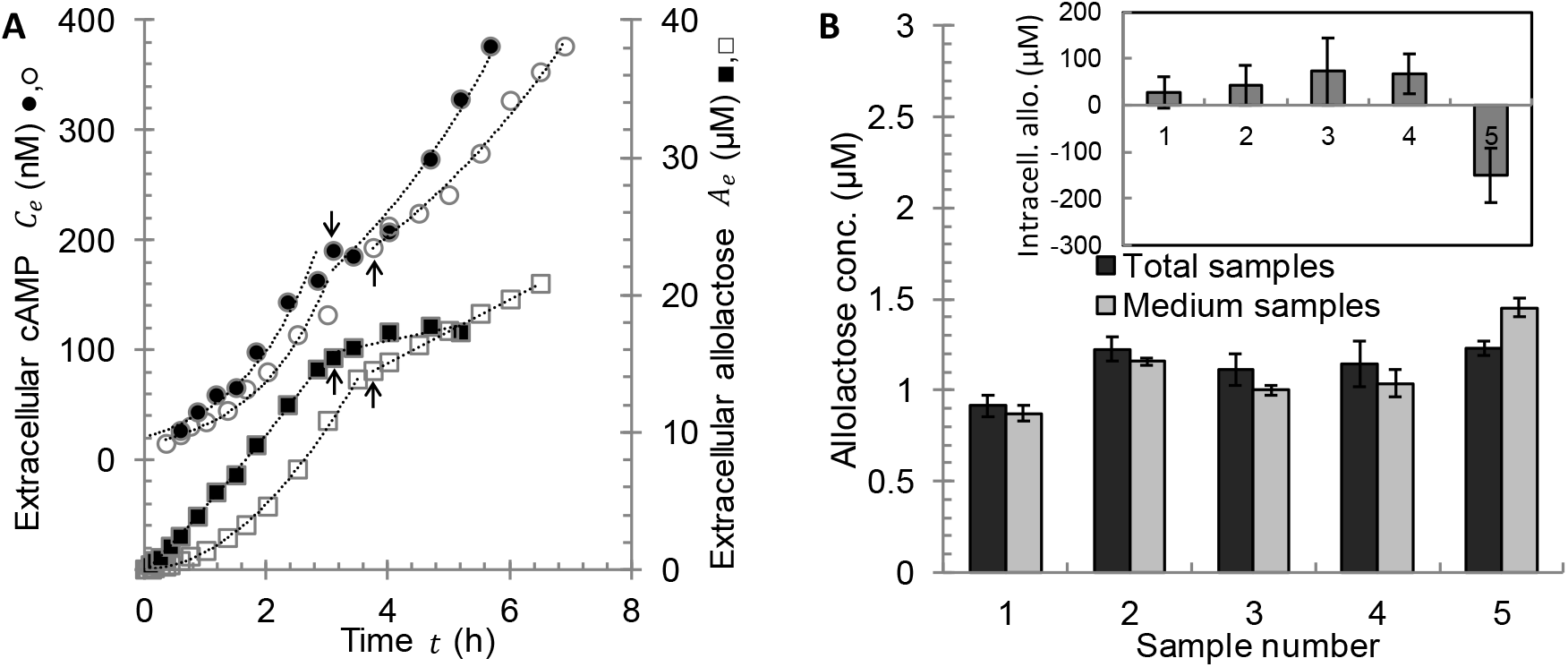
There is substantial efflux of allolactose and cAMP during growth of *E. coli* K12 MG1655 on lactose + maltose. (A) Evolution of extracellular cAMP (circles) and allolactose (squares) concentrations after inoculation of two shake flasks, each containing 4 mM lactose and 2.8 mM maltose, with uninduced cells grown on maltose (open symbols) and pre-induced cells grown on lactose + maltose (closed symbols). The arrows show the times at which some of the cells were removed in order to decrease the cell density. (B) The difference between the mean allolactose concentrations in total and medium samples, based on three samples, is on the order of the measurement error. The allolactose concentrations in the total and medium samples were measured 6 minutes after inoculation in five independent experiments of the type described in (A). The inset shows that the intracellular concentrations calculated by the difference method suffer from unacceptably high error. In this calculation, the cellular water volume was assumed to be 2.7 ml gdw^−1^ (48).

### 2.3 Measurement of β-galactosidase activity

The specific enzyme activity of β-galactosidase was assayed by Miller’s method (38).

### 2.4 Measurement of sugar concentration

To quantify sugar concentrations, we used high-performance anion-exchange chromatography with pulsed amperometric detection (HPAEC-PAD) in a Dionex ICS3000 system (Dionex Corp., Sunnyvale, CA, USA) (40). Sugars were separated at 30 °C on a CarboPac PA1 column (4 mm× 250 mm) connected to a CarboPac PA1 guard column (Dionex) using eluent containing 10 mM NaOH and 2 mM Barium Acetate (Eluent A) to precipitate the bicarbonate (41). Appropriately diluted samples were freshly thawed and sparged with N_2_ for >45 s. Manual injections (100 μl loop) were used to ensure reproducibility and minimize sample usage. Peaks obtained were integrated using Chromeleon 6.8 software.

### 2.5 Measurement of cAMP concentration

The Dionex ICS3000 system was also used for measuring the extracellular cAMP concentrations. The chromatographic separation and quantification method were adapted from Bhattacharya et al (42). High-performance anion-exchange chromatography (HPAEC) with variable wavelength detector (VWD) was used. Separation of cAMP was carried out at 30 °C on IonPac AS11-HC column (4 mm × 250 mm) connected to IonPac AS11 guard column (Dionex) using gradient elution with a flow rate of 1.0 ml/min. Combinations of 100 mM sodium hydroxide (Eluent B) and deionized water (Eluent C) were used to create the gradients. Initially, 7% B and 93% C were mixed (7 mM NaOH) and maintained for 10 minutes. The gradient of B was increased linearly in 2 minutes to 50% B and 50% C (50 mM NaOH) and maintained for 6 minutes. This was followed by restoring the eluent concentration to 7 mM in 2 minutes. The column was re-stabilized with initial elution conditions for 5–8 minutes before the next injection. Samples were freshly thawed and sparged with N_2_ for >45 s.

### 2.6 Generation of induction curve for IPTG

LacY^−^ mutants were isolated using UV mutagenesis and tested for the absence of LacY activity by measuring the accumulation of ^14^C-Methyl-β-D-thiogalactoside (TMG) (38). When exposed to ^14^C-TMG, these mutants developed equal concentrations on either side of the cellular membrane (15). To generate the induction curve for IPTG, the LacY^−^ cells were grown for at least 10 generations on a medium containing maltose (11 mM) or glycerol (43 mM), and various levels of IPTG, after which the specific β-galactosidase activity was measured by Miller’s method.

### 2.7 Determination of growth, enzyme synthesis, and metabolite efflux kinetics

We are interested in the variation of the specific rates of growth *μ*, β-galactosidase synthesis 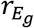, cAMP efflux 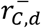 and allolactose efflux 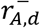 during the course of *lac* induction and repression. These rates were obtained by measuring the evolution of the concentrations of biomass *X*(*t*), β-galactosidase *E_g_*(*t*), extracellular cAMP *C_e_*(*t*), and extracellular allolactose *A_e_*(*t*), and then appealing to the following mass balances describing their evolution:

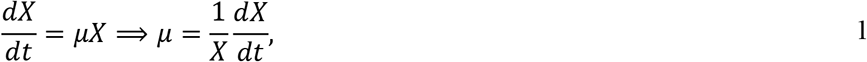

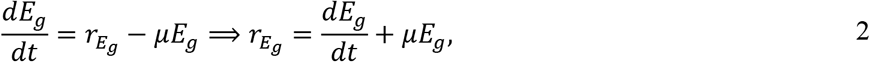

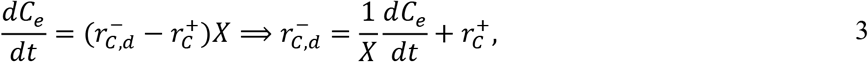

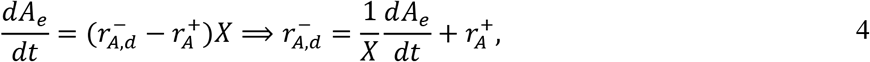

where *μE_g_* denotes the rate of β-galactosidase dilution (by growth), and 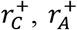 denote the specific rates of influx of intracellular cAMP and allolactose, respectively. Since we measured *X*(*t*), *E_g_*(*t*), *C_e_*(*t*), and *A_e_*(*t*), we could calculate *μ*, *dE_g_*/*dt*, and the net specific efflux rates of cAMP (1/*X*)(*dC_e_*/*dt*) and allolactose (1/*X*)(*dA_e_*/*dt*). Now, given *μ*, *E_g_*, and *dE_g_*/*dt*, we calculated the dilution rate *μE_g_*, and hence, the induction rate 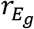 by appealing to Eq. (2). Also, given (1/*X*)(*dC_e_*/*dt*) and (1/*X*)(*dA_e_*/*dt*), we calculated the efflux rates 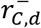 and 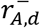 from Eqs. (3)–(4) provided the uptake rates 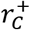 and 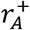 are known. We show below that there is no cAMP uptake 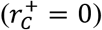, and the allolactose uptake rate 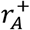 can be determined from suitable experiments.

We assume that cAMP and allolactose are expelled from the cell by diffusion, and their extracellular concentrations *C_e_*, *A_e_* are negligible compared to the corresponding intracellular concentrations denoted *C*, *A*, i.e.,

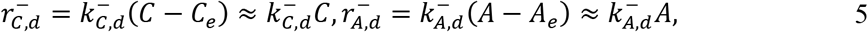

where 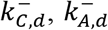 denote the diffusivities of cAMP and allolactose across the cell membrane. Epstein and co-workers showed that the specific efflux rate of cAMP was proportional to its intracellular concentration (10). We show below that the same is also true for allolactose.

### 2.8 Software

Data were processed using Microsoft Office Excel 2007 and MATLAB 2016b (The Mathworks, Inc.). In particular, the transient data were fitted by applying the least squares approximation to cubic B-spline fits obtained with *splinetool* in MATLAB.

## 3. Results

### 3.1 Measurement of intracellular allolactose by direct methods is prone to error

Since the cell-separation method involves the use of quenching agents that invariably cause metabolite leakage (31), we devised a thermal quenching method by isolating a mutant that expressed temperature-sensitive β-galactosidase. When these mutants were diluted in pre-heated medium at 60 °C, β-galactosidase was completely inactivated within two seconds (Fig. S3A). However, the allolactose concentrations, measured after filtering the quenched cells and extracting their intracellular allolactose in ~20 s, were extremely variable. This was probably due to rapid efflux of intracellular allolactose because when wild-type cells pre-loaded with [^14^C]TMG were diluted into pre-warmed medium 60 °C, 50 % of the [^14^C]TMG was lost within 20 seconds (Fig. S3B).

The efflux of intracellular allolactose also precluded the difference method. Indeed, when induced or non-induced cells of *E. coli* were grown in the presence of maltose + lactose, allolactose accumulated in the medium (Fig. 1A). Although the extracellular allolactose concentrations were not especially large (≤ 20 μM), the intracellular concentrations could not be determined precisely by the difference method. Fig. 1B shows that even when the mean concentration of extracellular allolactose was only ~1 μM, the mean concentration of total allolactose (in medium + cells) exceeded the mean concentration of extracellular allolactose (in medium) by ≲ 0.1 μM, which was comparable to the error of the measurements. The difference method is therefore susceptible to significant error because it involves calculation of the difference between two measured, and hence, error-prone, quantities of almost equal magnitudes, namely the total and extracellular allolactose concentrations.

### 3.2 There is significant consumption of extracellular allolactose, but not cAMP

The inherent errors of both the direct methods led us to consider an *indirect* method involving measurement of the specific allolactose efflux rate 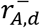 as a surrogate for the intracellular allolactose concentration. If the cells do not consume the extracellular allolactose, Eq. (4) implies that 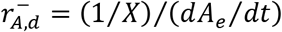 can be easily determined by measuring the concentrations of biomass and extracellular allolactose. However, Fig. 1A suggests that there was significant consumption of extracellular allolactose, but not cAMP, since the extracellular cAMP levels increase exponentially, whereas the extracellular allolactose levels increase rather slowly. We show below that this becomes more transparent if we plot the instantaneous concentrations of extracellular cAMP and allolactose in Fig. 1A against the corresponding biomass concentration. The slope of these *C_e_*(*t*) vs. *X*(*t*) and *A_e_*(*t*) vs. *X*(*t*) phase plots, commonly referred to as *differential plots*, are given by the relations

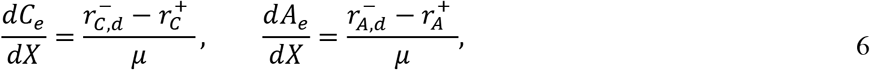

obtained by dividing Eqs. (3) and (4) by Eq. (1).

The shape of the differential plots for *pre-induced* cells immediately reveals if uptake of extracellular cAMP and allolactose is significant. To see this, observe that since pre-induced cells have been growing exponentially for several generations, they undergo balanced growth, so that 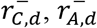, and *μ* are constant throughout the experiment (43). Now initially, 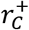 and 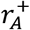 are zero because there is no cAMP and allolactose in the medium, and the slopes of the differential plots are 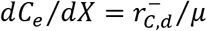 and 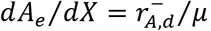, respectively. However, later on, 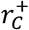 and 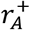 may become significant due to accumulation of cAMP and allolactose in the medium. We can determine if this is in fact the case by observing the subsequent values of the slope of the differential plot because it follows from Eq. (6) that the slope of the corresponding differential plot remains constant at its initial value if uptake is negligible compared to efflux throughout the experiment, and declines significantly from its initial value if uptake becomes comparable to expulsion during the course of the experiment. Fig. 2 shows that in pre-induced cells, the slope of the differential plot for cAMP remains constant (closed circles), which implies that cAMP uptake is negligible 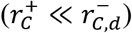 throughout the experiment. In contrast, the slope of the differential plot for allolactose decreases markedly and approaches zero at the end of the experiment (closed squares). It follows that there is significant allolactose uptake during the experiment, and the uptake rate almost equals the expulsion rate 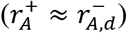 at the end of the experiment.

**Fig. 2.**
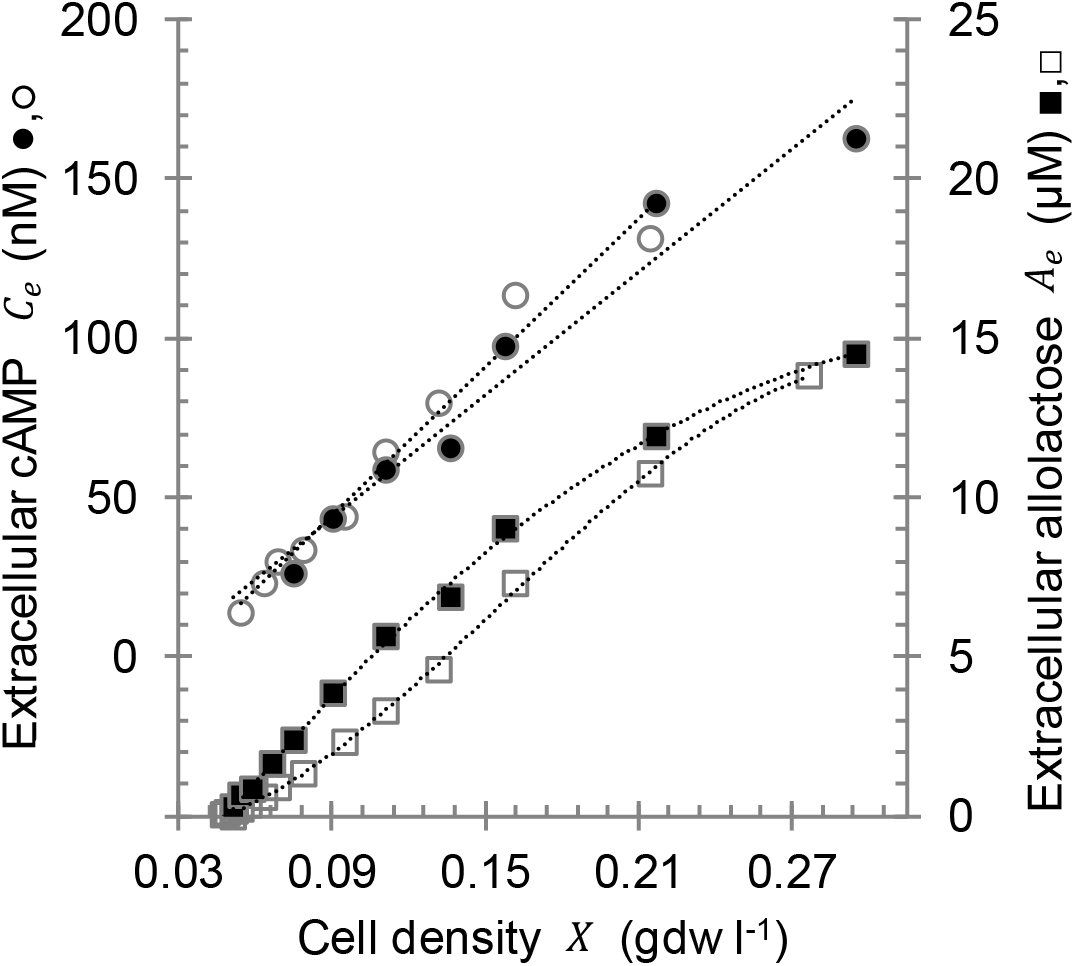
There is substantial uptake of allolactose, but not cAMP, during growth of *E. coli* K12 MG1655 on maltose + lactose. Differential plots of the data in Fig. 1A showing the variation of the extracellular cAMP (circles) and allolactose (squares) concentrations with the cell density in cultures inoculated with pre-induced (closed symbols) and uninduced (open symbols) cells. The differential plots for cAMP are linear throughout the experiment, and essentially the same slope is obtained with both pre-induced (●) and uninduced cells (○). However, the slopes of the differential plots for allolactose declines substantially towards the end. In pre-induced cells (■), the slope declines monotonically, whereas in uninduced cells (□), the slope increases and then decreases, with an inflection point at *X ≈* 0.12 gdw l^−1^.

The slopes of the differential plots for cAMP during the growth of pre-induced and non-induced cells (closed and open circles, resp. of Fig. 2) are essentially the same, which suggests that non-induced cells also do not consume cAMP 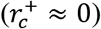. It follows from Eq. (3) that the specific cAMP efflux rate, a surrogate for the intracellular cAMP level in both steady state and transient conditions (10), is given by the relation

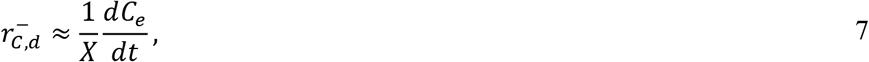

which can be calculated from the biomass and extracellular cAMP concentrations. Henceforth, we shall only be concerned with the specific allolactose efflux rate since our goal is to explore the existence of the allolactose-mediated positive feedback loop.

### 3.3 Development of a method for quantifying the specific allolactose uptake rate

Since there is significant allolactose uptake, Eq. (4) implies that the specific allolactose efflux rate 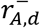 cannot be determined from the concentrations of biomass and extracellular allolactose unless the specific allolactose uptake rate 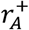 is known. We found that allolactose uptake was mediated primarily by Lac permease because 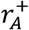 decreased 30-fold in *lacY^−^* mutants (Section S2). We show below that the kinetics of allolactose uptake in pre-induced cells can be quantified experimentally, and these kinetics allow us to estimate 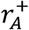 in cells induced to any level.

We determined the kinetics of permease-mediated allolactose uptake in pre-induced cells by analyzing the data in Fig. 1A. Since these cells are in a physiological steady state, they expel allolactose at a constant rate 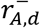, thus resulting in a progressive increase of the concentration *A_e_*(*t*) and specific uptake rate 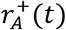 of extracellular allolactose, which can be determined as described below. Upon plotting 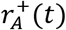 against *A_e_*(*t*), we obtain the kinetics of allolactose uptake in pre-induced cells.

We determined *A_e_*(*t*) and 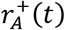 as follows. The concentration of extracellular allolactose was measured at various times (closed squares in Fig. 3A), and *A_e_*(*t*) was determined by fitting the data to cubic B-splines. Since we also measured the biomass concentration *X*(*t*) at various times, we could also determine the evolution of the net specific allolactose efflux rate (1/*X)(dA_e_*/*dt)* (open squares in Fig. 3A). The latter immediately yields the specific allolactose uptake rate 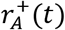 of pre-induced cells. To see this, observe that since 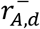 is constant in such cells, Eq. (4) can be rewritten as

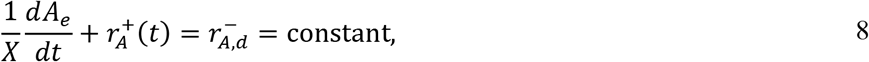

which implies that the increment of 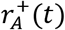 must equal the decline of (1/*X)(dA_e_*/*dt)*, i.e.,

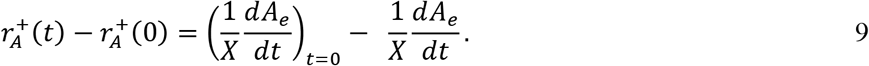

**Fig. 3:**
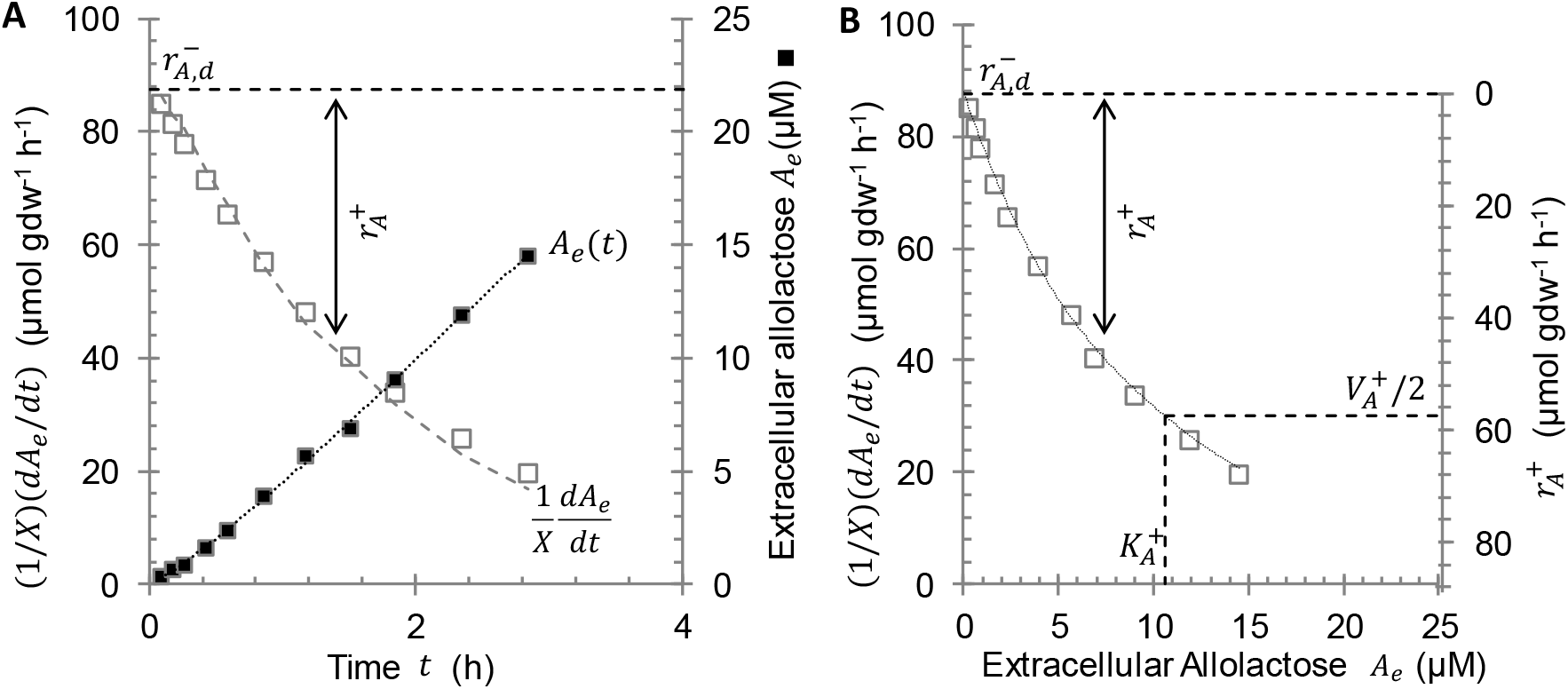
The specific allolactose uptake rate 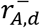 of pre-induced cells of *E. coli* K12 MG1655 increases hyperbolically with the extracellular allolactose concentration. (A) The net specific allolactose uptake rate 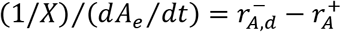 (□) was determined from the measured concentrations of biomass *X* (not shown) and extracellular allolactose *A_e_* (■). Since 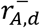 constanst in pre-induced cells, the specific allolactose uptake rate 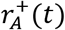 equals the decline of (1/*X*)/(*dA_e_*/*dt*) from its initial value 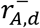, as as indicated by the double-headed arrow. (B) Determination of the kinetic parameters for allolactose uptake and efflux by pre-induced cells. The open squares are obtained by plotting the instantaneous values of (1/*X*)/(*dA_e_*/*dt*) in (A) against the corresponding values of *A_e_*(*t*). The solid curve shows the graph obtained when these data are fitted to Eq. (12). The dashed lines show the best-fit parameter values of 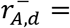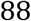 μmol gdw^−1^ h^−1^, 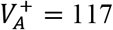 μmol gdw^−1^ h^−1^ and 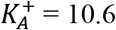 μM.

Now, since there is no allolactose in the medium initially, there is no allolactose uptake, i.e., 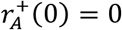, and the net specific allolactose efflux rate reflects pure efflux, i.e., 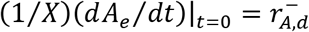. Hence, it follows from Eq. (9) that

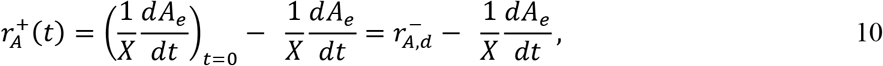

which shows that in pre-induced cells, 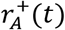 equals the decline of the measurable quantity (1/*X*)(*dA_e_*/*dt*) from its initial value 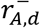 (arrow in Fig. 3A).

Fig. 3B shows the graph obtained when the instantaneous values of (1/*X*)(*dA_e_*/*dt*) in Fig. 3A are plotted against the corresponding values of *A_e_*(*t*). Since 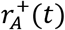 equals the decline of the measurable quantity (1/*X*)(*dA_e_*/*dt*) from its initial value 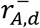, the graph suggests that 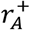 increases hyperbolically with the extracellular allolactose concentration *A_e_*. We confirmed this by determining the fit of the (1/*X*)(*dA_e_*/*dt*) vs *A_e_* data obtained under the assumption

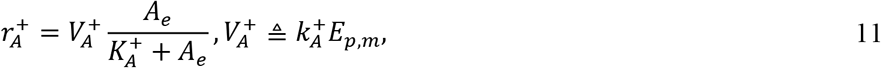

where *E_p,m_* denotes the (constant) specific permease activity of pre-induced cells. Then Eq. (10) can be rewritten as

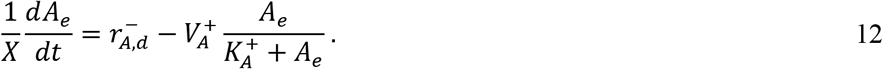

Upon fitting the (1/*X*)(*dA_e_*/*dt*) vs *A_e_* data (Fig. 3B) to Eq. (12), we obtained a good fit with the parameter values 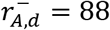 μmol gdw^−1^ h^−1^, 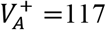 μmol gdw^−1^ h^−1^, and 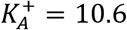 μM. Interestingly, the saturation constant for allolactose uptake 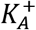, which was obtained during growth on a mixture of lactose (4 mM) and maltose, is only 4 % of the saturation constant for lactose uptake 270 μM during growth on pure lactose (44). Thus, LacY displays a remarkably high affinity for allolactose even in the presence of 4 mM lactose. We shall return to this point in the Discussion.

Given the kinetic parameters, 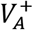 and 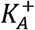, for allolactose uptake in pre-induced cells, we can estimate the specific allolactose uptake rate in partially induced cells as well. To see this, observe that allolactose uptake is mediated by the permease, and pre-induced cells follow the hyperbolic kinetics given by Eq. (11). It is therefore reasonable to assume that in partially induced cells, the specific allolactose uptake rate is proportional to the prevailing specific permease activity *E_p_*, i.e.,

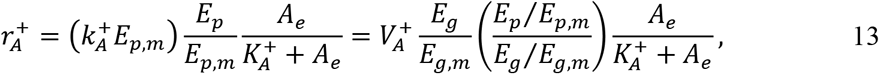

where *E_g_* and *E_g,m_* denote the specific β-galactosidase activity of partially and fully induced cells, respectively. In the prequel to this work (15), we have shown that in the absence of glucose, *E_p_* is proportional to *E_g_* due to coordinate expression of *lacY* and *lacZ*, and hence (*E_p_*/*E_p,m_*)/(*E_g_*/*E_g,m_*) = 1. However, in the presence of glucose, *E_p_* increases with *E_g_* faster than linearly because the more induced cells, the smaller the fraction of permease molecules inactivated by inducer exclusion. Hence, (*E_p_*/*E_p,m_*)/(*E_g_*/*E_g,m_*) is not constant, but an increasing function of *E_g_* which can be determined from the data in Fig. 2 of (15). Thus, if we measure *X*(*t*), *A_e_*(*t*), and *E_g_*(*t*) during *lac* induction in the presence of lactose + maltose or *lac* repression in the presence of lactose + glucose, we can calculate not only (1/*X*)(*dA_e_*/*dt*), but also 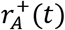 from Eq. (13), and substitute these quantities in Eq. (4) to obtain the specific allolactose efflux rate 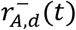, a surrogate for the intracellular allolactose concentration.

### 3.4 Intracellular allolactose levels and induction/dilution rates during *lac* induction

We are now ready to test the existence of the hypothesized allolactose-mediated positive feedback loop by determining if the intracellular allolactose concentration and induction rate increase with the induction level during the course of *lac* induction. To this end, we exposed non-induced cells to a mixture of lactose (4 mM) and maltose (2.8 mM), and measured the subsequent evolution of *X*(*t*), *A_e_*(*t*), and *E_g_*(*t*). These measurements allowed to calculate not only the specific allolactose efflux rate 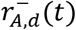 as described above, but also the induction and dilution rates, 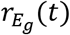, *μ*(*t*)*E_g_*(*t*) as described below. If 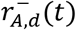 is proportional to the intracellular allolactose concentration, an assumption confirmed below, then the plots of 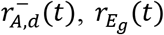, and *μ*(*t*)*E_g_*(*t*) against *E_g_*(*t*) yield the desired variation of the intracellular allolactose concentration and induction/dilution rates with the induction level.

The instantaneous specific allolactose efflux rates 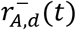 were estimated by two methods.

#### 1. Semi-empirical method

The net specific allolactose efflux rate (1/*X*)(*dA_e_*/*dt*) was first calculated from the *X* vs. *t* and *A_e_* vs. *t* data (open triangles and squares, resp. in Fig. S5A). Next, the specific allolactose uptake rate 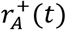 was estimated from the *A_e_* vs. *t* and *E_g_* vs. *t* data (open squares and circles, resp. in Fig. S5A) by using Eq. (12) with (*E_p_*/*E_p,m_*)/(*E_g_*/*E_g,m_*) = 1. Finally, the values of (1/*X*)(*dA_e_*/*dt*) and 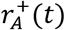 thus obtained, which are represented in Fig. S5B by *x* and *+*, respectively, were added, in accordance with Eq. (4), to determine the specific allolactose efflux rate 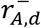 (open squares in Fig. S5B). We refer to this as the semi-empirical method for determining 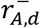 since Eq. (12) appeals to the assumption 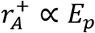, which is plausible, but not supported by direct evidence.

#### 2. Empirical method

To validate the above semi-empirical method, we also determined 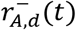 at various instants, denoted S1, S2, S3 in Fig. S5A, by the *initial rate method*. At these instants, some of the cells in the culture were transferred to a shake flask containing pre-warmed fresh medium and lactose (4 mM) + maltose (2.8 mM), but no allolactose. Thereafter, *X* and *A_e_* were measured for 10–20 min (closed triangles and squares, resp. in Fig. S5A). Under this condition, there is almost no allolactose uptake 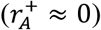, and the specific allolactose efflux rate is given by the expression 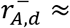 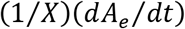. The specific allolactose efflux rates thus determined, which are represented in Fig. S5B by closed squares, were almost identical to those obtained by the semi-empirical method (open squares in Fig. S5B).

Since both methods yielded similar values of 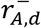, this quantity was estimated by the more convenient semi-empirical method in all subsequent experiments.

The instantaneous induction and dilution rates of β-galactosidase, 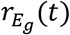 and *μ*(*t*)*E_g_*(*t*), were calculated as follows from the *X* vs. *t* and *E_g_* vs. *t* data in Fig. S5A. We first determined the instantaneous specific growth rate *μ*(*t*) = (1/*X*)/(*dX*/*dt*), β-galactosidase dilution rate *μ*(*t*)*E_g_*(*t*), and rate of change of the induction level *dE_g_*/*dt*. We then calculated 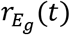 from the relation

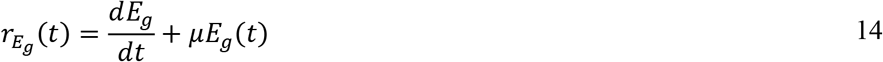

obtained from Eq. 2. The induction and dilution rates thus obtained are shown in Fig. S5C.

Fig. 4A shows the graph obtained upon plotting the specific efflux rate 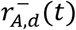 in Fig. S5B against the corresponding induction level *E_g_*(*t*) in Fig. S5A It can be seen that 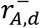, which is shown later on to be proportional to *A*, increases in a biphasic manner. Initially, it increases rapidly while *E_g_* is essentially constant, but thereafter 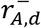 and *E_g_* increase simultaneously. The rapid initial increase is presumably due to the burst of allolactose produced immediately after lactose enters the uninduced cells. After this initial transient relaxes, the intracellular allolactose level, which is now in quasi-steady state, *increases* with the induction level. These data contradict the claim that the quasi-steady state intracellular allolactose concentration is *independent* of the induction level (23–25). Instead, they support the hypothesis that lactose enzymes promote the accumulation of intracellular allolactose. Since intracellular allolactose is known to stimulate synthesis of the lactose enzymes (2), the potential for positive feedback clearly exists. However, it remains to confirm that positive feedback does exist, and to explain how it leads to fully induced cells.

**Fig. 4.**
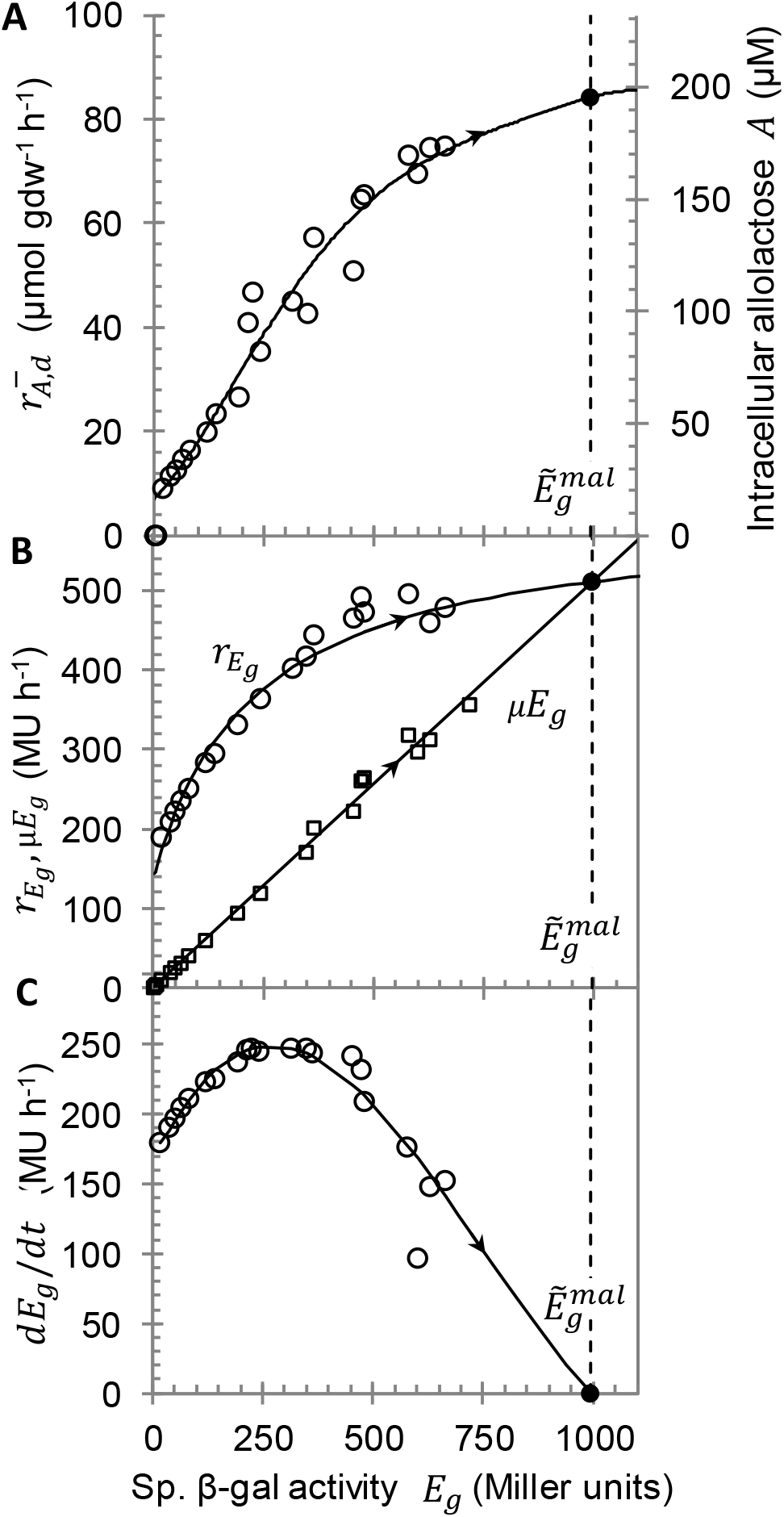
Positive feedback exists and drives induction by forward turning of the positive feedback loop. The phase plots are derived from the instantaneous *E_g_*(*t*) and 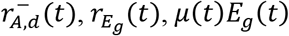 obtained during growth of uninduced cells of *E. coli* K12 MG1655 on maltose (2.8 mM) + lactose (4.0 mM) (Fig. S5). The arrows indicates the direction of evolution in time. (A) After an initial burst, the specific allolactose efflux rate 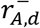 and the corresponding *A* (estimated from Fig. 6B) increase with *E_g_*, which implies that positive feedback can exist. (B) After an initial burst, the induction rate 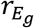 increases with *E_g_*, which shows that positive feedback does exist. The steady state induction level ^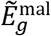^ (●) lies at the intersection of 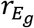 and *μE_g_*. (C) The net induction rate 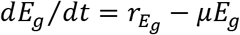 is positive for all 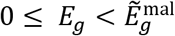.

We confirmed that positive feedback exists and leads to fully induced cells by plotting the induction and dilution rates, 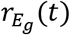 and *μ*(*t*)*E_g_*(*t*), in Fig. S5C against the corresponding induction level *E_g_*(*t*) in Fig. S5A (Fig. 4B). The plot shows that 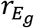 mirrors the biphasic profile of 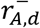. Initially, 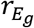 increases rapidly while *E_g_* is essentially constant, which reflects the initial burst of allolactose synthesis. This is followed by a subsequent slow phase during which 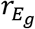, which is now in quasi-steady state, increases with *E_g_*. This increase provides direct evidence of positive feedback — the higher the induction level, the faster the induction rate. Moreover, since this positive feedback occurs in the absence of glucose, it is clear that inducer exclusion, which can be a source of positive feedback (Section S1.2.3), is not necessary for positive feedback. Fig. 4B also reveals how positive feedback drives the cells towards the fully induced steady state. Indeed, the steady state induction level, denoted 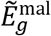, lies at the intersection of the 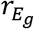 and *μE_g_* curves. Now, for all 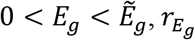 lies above *μE_g_*, so that 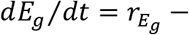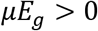 (Fig. 4C). Hence, the induction level of initially uninduced cells (*E_g_*(0) ≈ 0) increases, slowly but relentlessly, until it approaches the steady state level 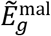. During this slow increase of the induction level, the quasi-steady state intracellular allolactose level increases as described by Fig. 4A. Thus, the quasi-steady concentrations both *E_g_* and *A* increase in tandem as predicted by our model.

### 3.5 Intracellular allolactose levels and induction/dilution rates during *lac* repression

We have also hypothesized that glucose-mediated catabolite repression occurs primarily because the forward turning of the positive feedback loop that leads to induction in the presence of lactose reverses direction upon the addition of glucose.

To test this hypothesis, we allowed uninduced cells to grow on maltose (2.8 mM) + lactose (4 mM) until the induction level reached steady state, at which point these fully induced cells were transferred to a medium containing glucose (2.2 mM) + lactose (4 mM) (*t* = 0 in Fig. S6). We measured the evolution of *X*(*t*), *A_e_*(*t*), *E_g_*(*t*) throughout the experiment, and calculated the corresponding 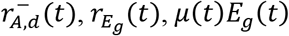, and *dE_g_*/*dt* by the methods described above. Upon plotting these calculated values against the corresponding values of *E_g_*(*t*), we obtained the graphs shown in Fig. 5 where the dashed and solid curves show, respectively, the fits to the data obtained during induction of the uninduced cells in the presence of maltose + lactose, and subsequent repression of the fully induced cells upon their transfer to glucose + lactose. The arrows in the figure indicate the direction of temporal evolution.

**Fig. 5:**
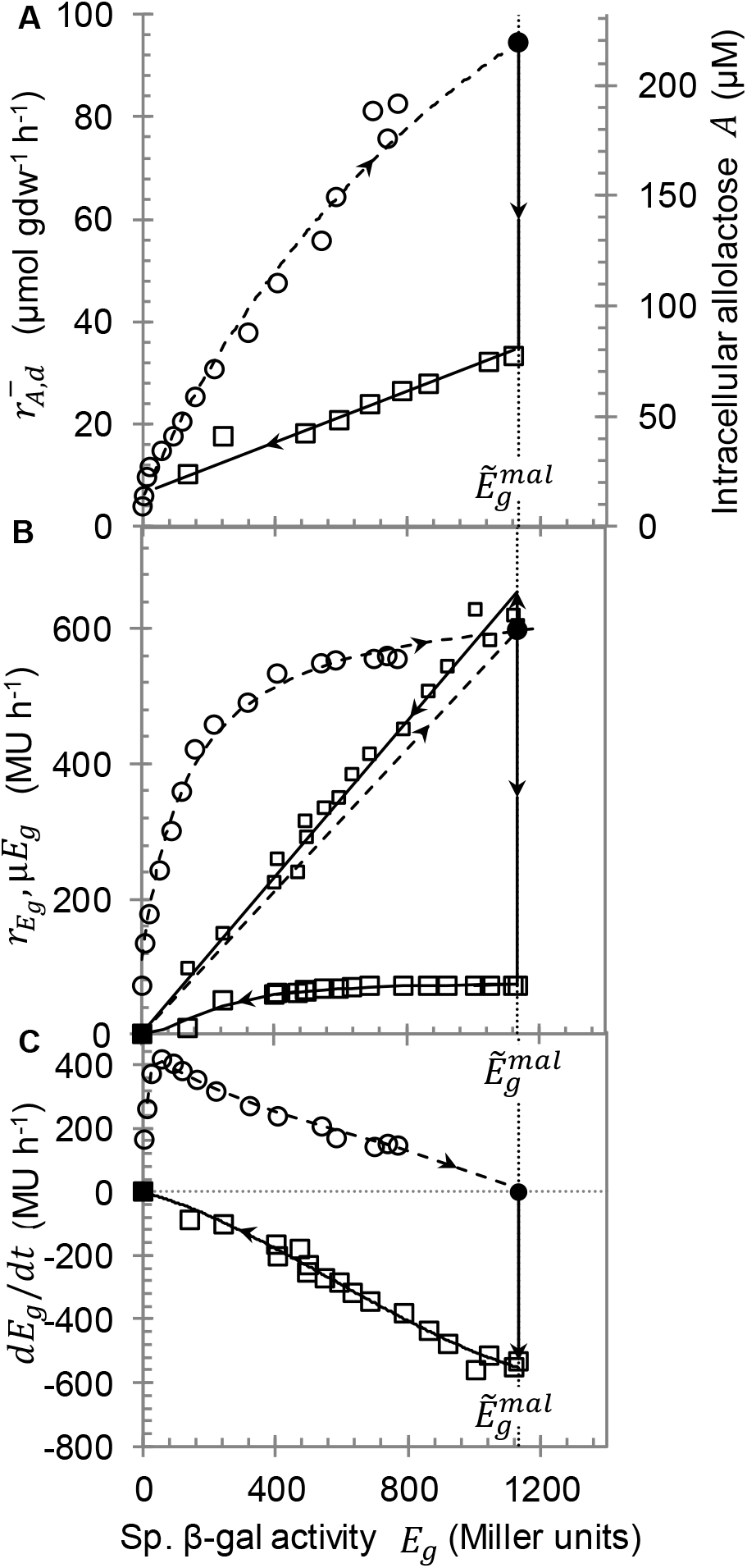
Positive feedback drives repression by reversed turning of the positive feedback loop. The dashed and solid phase plots are derived from the instantaneous *E_g_*(*t*) and 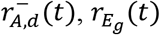, *μ*(*t*)*E_g_*(*t*) obtained during induction of uninduced cells of *E. coli* K12 MG1655 on maltose (2.8 mM) + lactose (4 mM), and repression of the induced cells thus obtained upon transfer to glucose (2.2 mM) + lactose (4 mM), respectively. (A) Upon transfer to glucose + lactose, the specific allolactose efflux rate 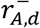 and the corresponding *A* (estimated from Fig. 6B) decrease abruptly at constant *E_g_*, which is followed by a gradual decline with *E_g_*. (B) Upon transfer to glucose + lactose, the dilution rate *μE_g_* remains unchanged, and the induction rate 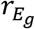 shifts downward ~6-fold, but the steady state induction level ^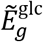^ (■) reduces 1000-fold. (C) Upon transfer to glucose + lactose, _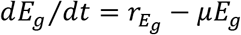_ is negative for all ^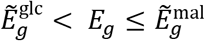^.

If 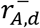 is a surrogate for *A*, then it is clear from the solid curve of Fig. 5A that immediately after the addition of glucose, *A* decreases 3-fold from its steady state value during exponential growth on lactose + maltose, whereas *E_g_* remains nearly constant. This rapid decline of *A* at constant *E_g_* is reminiscent of inducer exclusion: upon addition of glucose, permease, but not β-galactosidase, is partially inactivated, and *A* responds by instantly decreasing to a lower quasi-steady state level. After this rapid transient, the induction level *E_g_* decreases, and so does the quasi-steady state intracellular allolactose level *A* as it continuously adjusts to the slowly declining induction level *E_g_*. This concurrent decline of *A* and *E_g_* is consistent with the trend expected from a reversal of the positive feedback loop.

It remains to explain what drives the reversal of the positive feedback loop. This requires a closer look at the dynamics of the induction level in the presence of glucose + lactose, which are governed by the LacZ induction and dilution rates (solid curves in Fig. 5B). Evidently, positive feedback persists (i.e., induction follows autocatalytic kinetics) even in the presence of glucose since 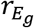 is an increasing function of *E_g_*. We show below that the positive feedback loop is reversed because these autocatalytic induction kinetics conspire with inducer exclusion and cAMP-mediated regulation to shift the steady state induction level to a new value 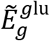 that is drastically lower than the steady state value 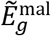 sustained in the presence of maltose + lactose. To see this, observe that after the fully induced cells are transferred to a medium containing glucose, the dilution rate remains essentially unchanged (compare dashed and full lines in Fig. 5B), but the induction rate shifts downward (compare dashed and full curves in Fig. 5B). Although the induction rate declines only 6-fold, the induction level 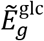 at the new steady state, which occurs at the intersection of the solid curves representing the induction and dilution rates (■), is 1000-fold lower than the induction level 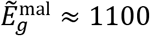 MU at the steady state before the addition of glucose (●). Moreover, for all 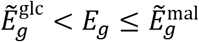, the dilution rate exceeds the induction rate, so that 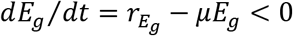 (solid curve of Fig. 5C). Hence, *E_g_* decreases, slowly but relentlessly, from its initial value 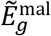 to the drastically lower new steady state value 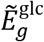. As the induction level declines slowly, so does the quasi-steady state intracellular allolactose level as described by the oblique solid line Fig. 5A. Thus, the reversal of the positive feedback loop is driven by the dramatic shift of the steady state induction level in the presence of glucose.

In the presence of gratuitous inducers, positive feedback exists (21, 22) and drives glucose-mediated catabolite repression due to the reversal of the positive feedback loop (15). Now, comparison of Fig. 5A–C with Fig. 5A–C of (15) shows that the kinetics observed upon addition of glucose to a culture growing on lactose are formally similar to those observed upon addition of glucose to a culture growing on glycerol and 100 μM TMG. It is therefore reasonable to conclude that the same mechanism, namely reversal of the positive feedback loop, drives glucose-mediated repression in the presence of both TMG and lactose.

In principle, the 6-fold decline of the induction rate immediately the addition of glucose (Fig. 5B) could be due to both inducer exclusion and cAMP-mediated regulation, but the data show that inducer exclusion plays no role initially because at this time, 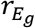 is saturated with respect to *A*. To see this, observe that even after the initial 3-fold decline of *A* (Fig. 5A), 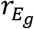 remains constant while *E_g_* decreases from 1100 to 500 MU (Fig. 5B), and hence, *A* decreases from 80 to 30 μM (Fig. 5A). It follows that the initial 6-fold decline of 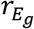 must be due to cAMP-mediated regulation, which is consistent with the cAMP-mediated *severe transient repression* that is known to strongly inhibit *lac* expression for ~1 doubling time after glucose addition, as opposed to the cAMP-mediated *weak permanent repression* that persists thereafter (4). However, later on, when *E_g_* has fallen below 500 MU, 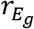 is unsaturated with respect to *A*, and inducer exclusion can also play a role in the decline of the induction rate.

### 3.6 The allolactose efflux rate is proportional to the intracellular allolactose level

Thus far, we have assumed that 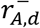 is proportional to A. We show here that this is indeed the case, and determine the corresponding proportionality constant.

To this end, we first determined the induction rate 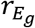 at various 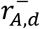 by calculating, as described above, the evolution of 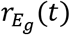 and 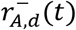 during the growth of uninduced cells on lactose (4 mM) + maltose (2.8 mM) (crosses and open squares in Fig. S7). It is useful to normalize 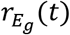 by its value in fully induced cells because the theory suggests (Section S3), and the data discussed below confirm, that this normalization eliminates the effect of cAMP on the induction rate. Upon plotting this normalized induction rate 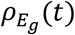 against 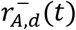, we obtained the 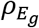 vs 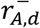 plot (open squares of Fig. 6A), which will be referred to as the normalized induction curve with respect to 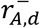. Since 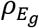 is independent of the cAMP level, we may assume that it is completely determined by the intracellular allolactose concentration *A*. As we show next, this property enables the estimation of *A* corresponding to any given 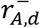.

**Fig. 6.**
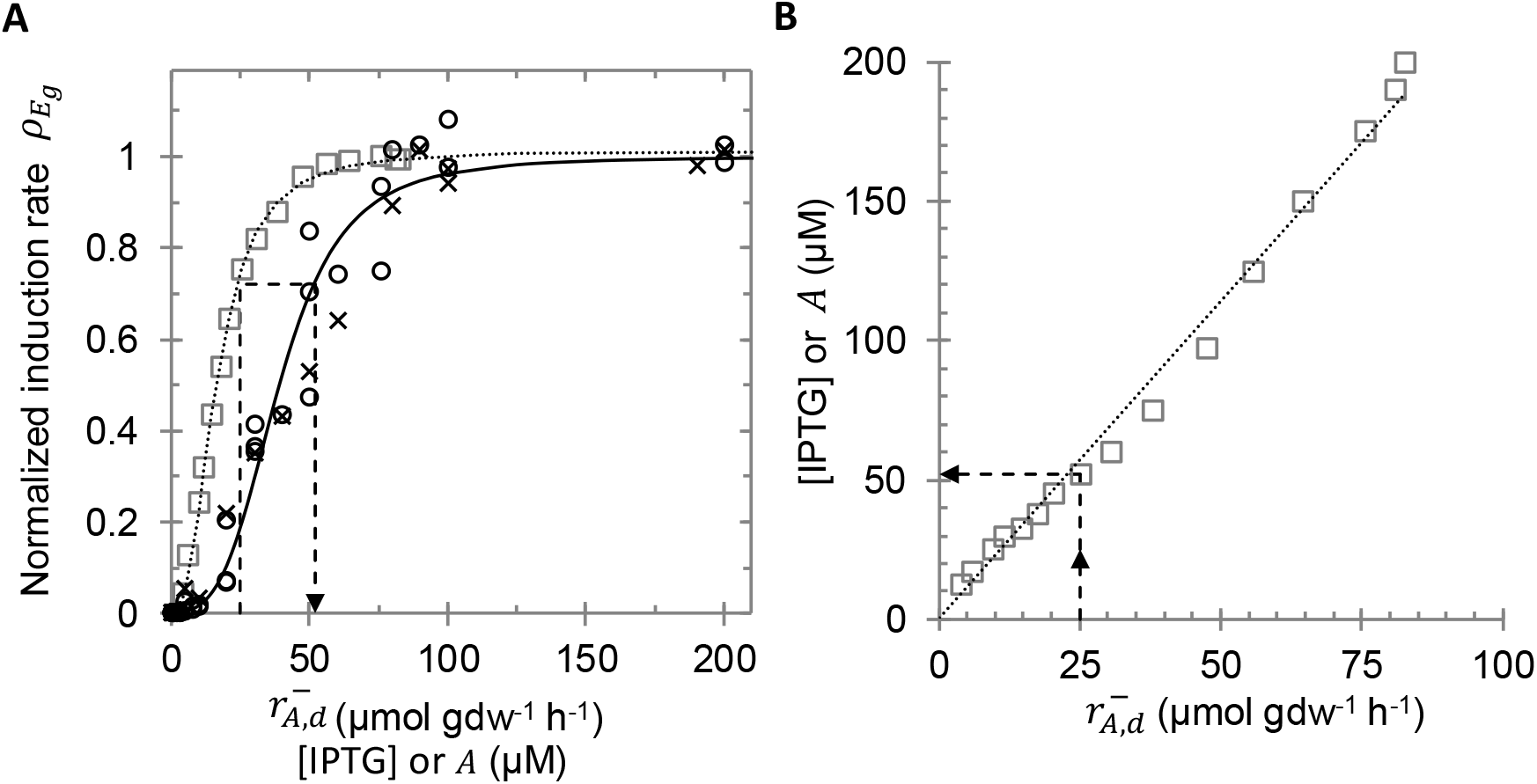
The intracellular concentration of allolactose *A* is proportional to its specific efflux rate 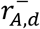. (A) The dashed curve and open squares show the variation of the normalized induction rate 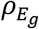 with 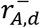, which is derived from the data obtained during growth of uninduced cells on maltose + lactose (Fig. S7). The solid curve shows the variation of 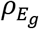 with the extracellular IPTG concentration [IPTG], which is derived from the steady state induction rates obtained in *lacY^−^* cells exposed to various concentrations of extracellular IPTG in the presence of glycerol (○) and maltose (*x*). Given any 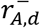, the dashed arrows indicate the algorithm for determining the corresponding value of [IPTG], and hence *A*. (B) The eaxlltorlaaccetlolsuelacroInPcTeGntroartiionntrsacoelrlruelsapronding to various value of 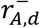. The solid line shows that the estimated intracellular allolactose concentration *A* is proportional to 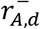.

To estimate *A* corresponding to any given 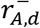, one must ideally determine the normalized induction curve with respect to allolactose (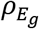 vs. *A*), which can be obtained by measuring and normalizing the induction rates obtained in *lacY^−^ lacZ^−^* cells exposed to a non-inducing carbon source, such as glycerol or maltose, and various concentrations of extracellular allolactose. Given the 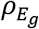 vs. 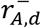 and 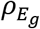 vs. A curves, one can calculate the value of A corresponding to any given 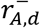 as indicated by the dashed lines in Fig. 6A. First, find the value of 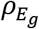 corresponding to the given 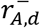 by using the 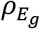 vs. 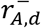 curve, and then find the value of a that yields this value of 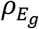 by using the 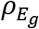 vs. curve. Thus, the value of corresponding to any given 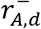 is that value of which provides the same normalized induction rate 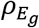 as that obtained from the given value of 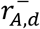.

We could not determine the 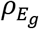 vs *A* curve because allolactose is not commercially available, but this curve is essentially identical to the normalized induction curve with respect to IPTG (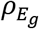 vs. [IPTG]), which can be easily obtained by measuring and normalizing the induction rates in *lacY^−^* cells exposed to a non-inducing carbon source and various IPTG concentrations. Indeed, the theory suggests that the 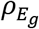 vs. *A* and 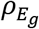 vs. [IPTG] curves coincide if the dissociation constants for binding of Lac repressor to allolactose and IPTG are the same (Section S3). The experiments confirm that allolactose and IPTG are indistinguishable as inducers since they bind to the repressor with the same dissociation constant of 0.1 μM and have the same effect on the repressor-operator complex (2). Hence, in the foregoing method for determining the value of *A* corresponding to 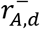, we can just as well use the 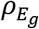 vs [IPTG], instead of the 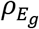 vs *A*, curve.

Fig. 6A shows the 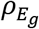 vs [IPTG] data obtained with two different non-inducing carbon sources, namely, maltose (*x*) and glycerol (○). At any given concentration of IPTG, the induction rates obtained with maltose and glycerol are quite different; in particular, under fully induced conditions, the induction rates were 768 and 1308 MU h^−1^, resp., which presumably reflect the different intracellular cAMP levels during growth on maltose and glycerol (10). However, the corresponding normalized induction rates in Fig. 6A are essentially the same, thus confirming that normalization eliminates the effect of cAMP.

The dashed lines of Fig. 6A describe the process by which we obtain the value of *A* corresponding to one particular value of 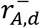. Application of this process to several values of 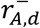 yields multiple pairs of the form 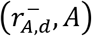, which were plotted to obtain the graph shown in Fig. 6B. It is clear that 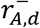 is essentially proportional to *A*, which implies that the specific allolactose efflux rate is a legitimate surrogate for the intracellular allolactose concentration.

## 4. Discussion

### 4.1 Critical appraisal of the data

The goal of this section is to assess the reliability of the data leading to the main conclusions stemming from this work:

1. Positive feedback exists and drives glucose-mediated repression.
2. Positive feedback is mediated by allolactose.

To this end, we recall the data leading to these conclusions, and examine the measurements and calculations leading to these data.

We reached the first conclusion by showing that during induction in the presence of lactose + maltose (Fig. 4B, C) as well as repression in the presence of lactose + glucose (Fig. 5B, C):

a. The quasi-steady state induction rate 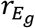 increases with the induction level *E_g_*.
b. The quasi-steady state induction and dilution rates in the presence of glucose + lactose are formally similar to those obtained in the presence of glucose + TMG, a condition in which the role of positive feedback in mediating strong repression has been demonstrated (15).

The quasi-steady state induction and dilution rates reported in these experiments were determined by measuring *X*(*t*) and *E_g_*(*t*), and calculating the instantaneous specific growth rate *μ(t) = (*1/*X)(dX*/*dt*), dilution rate *μ(t)E_g_(t*), and induction rate 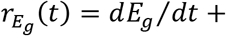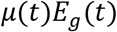. Since *X*(*t*) and *E_g_*(*t*) were measured using standard methods, and the calculations appealed to standard mass balances for *X* and *E_g_*, we can impute a high level of confidence to our first conclusion. More generally, our method for determining the induction and dilution rates from the data, which has not been reported before, enables systematic formulation of kinetic models of carbon catabolite repression (45, 46).

We arrived at the second conclusion by showing that during the course of induction (resp., repression), the quasi-steady state intracellular allolactose concentration *A*(*t*) and specific β-galactosidase activity *E_g_*(*t*) increase (resp., decrease) in tandem, thus concluding that *A* increases with *E_g_* (Fig. 4A, Fig. 5A). Here, *A* was estimated as follows:

a. We calculated the specific allolactose expulsion rate 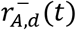 by two independent methods (Fig. S5), namely, the empirical method wherein we determined the initial net specific allolactose expulsion rate 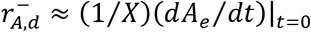 experimentally, and the semi-empirical method based on the mass balance 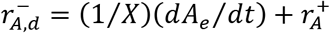, wherein we determined (1/*X*)(*dA_e_*/*dt*) experimentally and added to it the instantaneous value 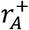 estimated from Eq. (13). Since both independent methods yielded almost identical values, our values of 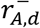 are quite reliable.
b. We estimated the value of *A* corresponding to any given value of 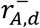 by determining the concentration of intracellular allolactose that gives the same normalized induction rate 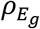 as that obtained from the given value of 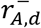 (Fig. 6). This method is based on two assumptions, namely allolactose and IPTG induce identically, and the normalized induction rate is completely determined by the intracellular inducer concentration. The first assumption is supported by the data in the literature (2), and the second assumption was validated by showing that in the presence of IPTG, the LacY^−^ strain yielded the normalized induction rates on maltose and glycerol.

We also measured the intracellular allolactose concentrations by an alternative technique aimed at reducing the error of the difference method. To this end, cells were grown on maltose + lactose in a perfusion reactor fed with high flow rates of fresh medium to reduce the extracellular allolactose concentration, and hence, the error of the difference method. The intracellular allolactose concentrations, obtained by applying the difference method to 10 samples of fully induced cells derived from the perfusion reactor, were 120 ± 100 μM (manuscript in preparation). The mean value is therefore comparable to our estimate of 200 μM (Fig. 6A), but the large error highlights the dire need for a high-precision method that overcomes the intrinsic errors of the existing direct methods.

It seems desirable to also compare our data with those in the literature. We are aware of only one study reporting intracellular allolactose concentrations. Huber et al. used the difference method to measure the intracellular allolactose concentrations in *lac*-constitutive cells exposed to 60 mM lactose (26). They reported intracellular allolactose concentrations of ~300 mM, which is much higher than our estimate of 200 μM in fully induced cells. This is probably due to the errors inherent in the difference method — Fig. 1 of their paper shows that the intracellular allolactose concentrations were frequently negative since the amount in the medium exceeded the total amount, which mirrors our data (Fig. 1B). In another study, Huber and co-workers also reported the efflux rates of glucose, galactose, and allolactose in *lac*-constitutive cells exposed to 1 mM lactose (47). They observed that almost all the lactose taken by the cells was rapidly expelled as glucose, galactose, and allolactose. We observed the same phenomenon with induced cells for the first 15 min (Table 1), but after this initial period, there was no net efflux of glucose and galactose (Fig. S8). We also find that the specific allolactose efflux rates in our induced cells are 15-fold less than those observed by Huber et al. This may be due to the 15-fold smaller lactose concentrations (4 mM) used in our experiments.

**Table 1.**
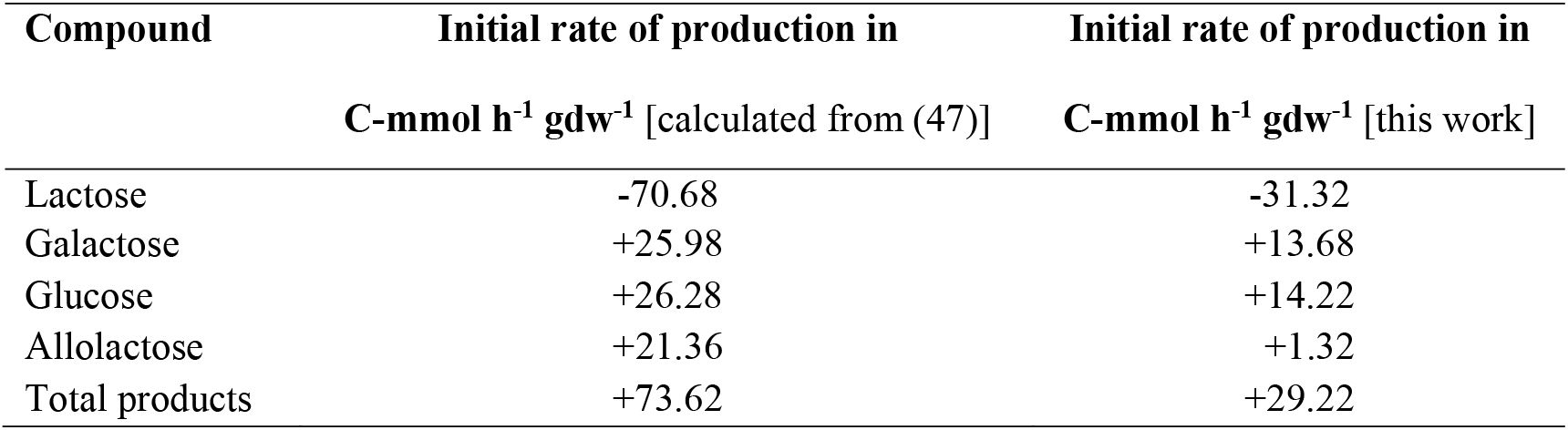
Comparison of initial rates of sugar production obtained in this study with the results of Huber et al. (1980). The negative rate of lactose production indicates consumption of lactose.

| Compound | Initial rate of production in | Initial rate of production in |
| --- | --- | --- |
|  | C-mm<sup>ol</sup> h<sup>-1</sup> gdw<sup>-1</sup> [calculated from (47)] | C-mm<sup>ol</sup> h<sup>-1</sup> gdw<sup>-1</sup> [this work] |
| Lactose | -70.68 | -31.32 |
| Galactose | +25.98 | +13.68 |
| Glucose | +26.28 | +14.22 |
| Allolactose | +21.36 | +1.32 |
| Total products | +73.62 | +29.22 |

### 4.2 Evolutionary implication of the high affinity of Lac permease for allolactose

It is remarkable that permease-mediated allolactose uptake occurs even in the presence of 4 mM lactose. Indeed, we found that the *apparent* saturation constant for permease-mediated allolactose uptake, observed in the presence of 4 mM lactose, was 10 μM, which is an order of magnitude smaller than the *intrinsic* saturation constant of 270 μM for permease-mediated lactose uptake (36, 48). It seems likely that the intrinsic saturation constant for permease-mediated allolactose uptake is even lower. If we assume simple competitive kinetics with lactose as the competitor, we find that the intrinsic saturation constant for allolactose uptake is 1 μM which is 250-fold smaller than the intrinsic saturation constant for lactose. This remarkably high affinity of Lac permease for allolactose suggests that the *lac* operon did not evolve for consumption of lactose. Egel made this claim earlier, but it was based on the observation that allolactose, rather than lactose, was the true inducer (49). Our data show that the permease has a much higher affinity for allolactose than lactose, which provides further evidence that the *lac* operon may have evolved to promote consumption of allolactose or a structurally similar compound.

### 4.3 Explaining the data adduced in support of the inducer exclusion hypothesis

In the Introduction, it was observed that several interventions that target inducer exclusion also abolish catabolite repression. Now we have shown above that positive feedback is the dominant mechanism for catabolite repression since inducer exclusion accounts only for the initial 3-fold decline of the allolactose level (as indicated by 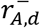), and the subsequent >30-fold decline is driven by the reversal of the positive feedback loop (Fig. 5A, B). Given this fact, it is reasonable to ask: If positive feedback, rather than inducer exclusion, is the dominant mechanism of catabolite repression, why do interventions that target inducer exclusion succeed in abolishing catabolite repression?

The answer to the above question might be that the interventions that targeted inducer exclusion abolished not only inducer exclusion, but also positive feedback. To see this, recall that positive feedback can occur only if the inducer stimulates lactose enzyme synthesis and the lactose enzymes promote inducer accumulation, i.e., the induction rate has the functional form 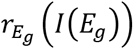 wherein 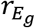 increases with the inducer concentration *I*, which in turn increases with *E_g_*. It follows that there is no positive feedback if

1. 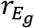 is independent of *I*, i.e., *lac* expression is independent of inducer levels as is the case in *lacI^−^* cells.
2. *I* is independent of *E_g_*, i.e., inducer accumulation is independent of lactose enzyme levels. This is the case when surplus IPTG is present in the medium since under these conditions, the main inducer is not allolactose, but IPTG which accumulates inside cells primarily by the LacY-independent mechanism of passive diffusion (50, 51).

In both cases, 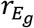 is independent of *E_g_*, i.e., *lac* expression is constitutive. Thus, the foregoing interventions targeting inducer exclusion also abolish positive feedback, which might explain their efficacy in abolishing catabolite repression.

## 5. Conclusions

In this work, we addressed a controversial debate on the mechanism underlying the strong >500-fold *lac* repression observed when glucose is added to a culture growing in the presence of lactose. Specifically, we have proposed that the allolactose-mediated positive feedback loop amplifies the small effects of inducer exclusion and cAMP-mediated transcriptional repression, but mathematical models show that the proposed positive feedback loop does not exist since the allolactose concentration is independent of the induction level. Thus, the two key questions of this debate are: Does the allolactose level increase with the induction level? Does positive feedback exist, and does it drive the repression?

To address these questions, we first showed the proposed positive feedback loop can exist in theory because the foregoing mathematical models are based on two assumptions — no lactose/allolactose expulsion and constant LacY:LacZ ratio — that are inconsistent with the data, and relaxing them restores positive feedback. We then showed that the proposed positive feedback loop does exist and drives the repression by performing two sets of experiments: an induction experiment in which uninduced cells were allowed to become fully induced during growth on maltose + lactose (Fig. 4), and a repression experiment in which the fully induced cells thus obtained were transferred to medium containing glucose + lactose (Fig. 5). The first question was resolved by developing a method for estimating the intracellular allolactose levels (Fig. 6). With the help of this method, we showed that in both experiments, the allolactose levels increase with the LacZ level, which implies that positive feedback *can* exist. The second question was resolved by constructing the LacZ induction and dilution rates from the data, a method developed in the prequel to this work. This method enabled us to show that in both experiments, the LacZ induction rate increases with the LacZ level, which implies that positive feedback *does* exist in the presence of lactose. The method also shows explains how positive feedback drives massive repression — when fully induced cells are exposed to glucose + lactose, the induction rate reduces only 6-fold, but the steady state induction level reduces 1000-fold, which forces the positive feedback loop to turn in the reverse direction.

Our data show that it not just the molecular mechanisms, but also systemic dynamical properties of the regulatory network, such as positive feedback, that drive glucose-mediated *lac* repression.

## Acknowledgement

We dedicate this work to late Prof. Frederick C. Neidhardt whose questions led to the experiments reported here. The early part of this research was supported by the DST grant SR/SO/BB-79/2010.

## ABBREVIATIONS AND NOTATIONS

### ABBREVIATIONS

cAMP: 3’,5’-cyclic adenosine monophosphate
CGSC: Coli Genetic Stock Centre, Yale University
gdw: Grams of cell dry weight, g
HPAEC: High-performance anion exchange chromatography (also see PAD)
IPTG: Isopropyl β-D-thiogalactopyranoside or MeSH
LB: Luria-Bertani broth
MU: Miller units
OD_600_: Optical density at 600 nm
ONPG: *ortho*-nitrophenol-β-D-galactopyranoside
PAD: Pulsed amperometric detection (also see HPAEC)
PES: Polyethersulfone membrane
TMG: Methyl-β-D-1-thiogalactopyranoside
[^14^C]TMG: Carbon-14 labelled isotope of TMG
UV: Ultraviolet (radiation)
VWD: Variable wavelength detector

### NOTATIONS

*A*: Intracellular allolactose concentration, μM
*A_e_*: Extracellular allolactose concentration, μM
*C*: Intracellular cAMP concentration, nM
*C_e_*: Extracellular cAMP concentration, nM
*E_g_*: Specific β-galactosidase activity, Miller units
*E_g,m_*: Specific β-galactosidase activity of pre-induced cells, Miller units
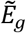: Steady state specific β-galactosidase activity, Miller units
*E_p_*: Specific Lac permease activity
*E_p,m_*: Specific Lac permease activity of pre-induced cells
*I*: Inducer concentration, μM
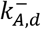: Diffusivity of allolactose through cell membrane, h^−1^
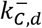: Diffusivity of cAMP through cell membrane, h^−1^
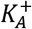: Apparent saturation constant for permease-mediated allolactose uptake, μM
OD_600_: Optical density at 600 nm
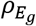: Normalized induction rate
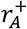: Specific allolactose influx rate into the cells, μmol h^−1^ gdw^−1^
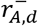: Specific allolactose efflux rate out of the cells, μmol h^−1^ gdw^−1^
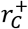: Specific cAMP influx rate into the cells, nmol h^−1^ gdw^−1^
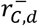: Specific cAMP efflux rate out of the cells, nmol h^−1^ gdw^−1^
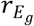: Specific synthesis rate of β-galactosidase, Miller units h^−1^
*t*: Time, s or min or h
*μ*: Specific growth rate of *E. coli* cells, h^−1^
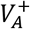: Maximum specific allolactose uptake rate, μmol h^−1^ gdw^−1^
*X*: Biomass or cell density, gdw l^−1^

## REFERENCES

1. Makman, R.S., and E.W. Sutherland. 1965. Adenosine 3’,5’-phosphate in *Escherichia coli*. J. Biol. Chem. 240: 1309–14.

2. Jobe, A., and S. Bourgeois. 1972. *lac* repressor-operator interaction. VI. The natural inducer of the *lac* operon. J. Mol. Biol. 69: 397–408.

3. Perlman, R.L., and I. Pastan. 1968. Cyclic 3′5′-AMP: Stimulation of β-galactosidase and tryptophanase induction in *E. coli*. Biochem. Biophys. Res. Commun. 30: 656–664.

4. Magasanik, B. 1970. Glucose effects: inducer exclusion and repression. In: The Lactose Operon. . pp. 189–219.

5. Loomis Jr., W.F., and B. Magasanik. 1964. The relation of catabolite repression to the induction system for β-galactosidase in *Escherichia coli*. J. Mol. Biol. 8: 417–426.

6. Hogema, B.M., J.C. Arents, R. Bader, K. Eijkemans, T. Inada, H. Aiba, and P.W. Postma. 1998. Inducer exclusion by glucose 6-phosphate in *Escherichia coli*. Mol. Microbiol. 28: 755–65.

7. Deutscher, J., C. Francke, and P.W. Postma. 2006. How Phosphotransferase System-Related Protein Phosphorylation Regulates Carbohydrate Metabolism in Bacteria. Microbiol. Mol. Biol. Rev. 70: 939–1031.

8. Inada, T., K. Kimata, and H. Aiba. 1996. Mechanism responsible for glucose-lactose diauxie in *Escherichia coli*: challenge to the cAMP model. Genes to Cells. 1: 293–301.

9. Narang, A. 2009. Quantitative effect and regulatory function of cyclic adenosine 5’-phosphate in *Escherichia coli*. J. Biosci. 34: 445–63.

10. Epstein, W., L.B. Rothman-Denes, and J. Hesse. 1975. Adenosine 3’:5’-cyclic monophosphate as mediator of catabolite repression in *Escherichia coli*. Proc. Natl. Acad. Sci. U. S. A. 72: 2300–4.

11. Narang, A. 2009. cAMP does not have an important role in carbon catabolite repression of the *Escherichia coli lac* operon. Nat. Rev. Microbiol. 7: 250.

12. Stülke, J., and W. Hillen. 1999. Carbon catabolite repression in bacteria. Curr. Opin. Microbiol. 2: 195–201.

13. Görke, B., and J. Stülke. 2008. Carbon catabolite repression in bacteria: Many ways to make the most out of nutrients. Nat. Rev. Microbiol. 6: 613–624.

14. Winkler, H.H., and T.H. Wilson. 1967. Inhibition of β-galactoside transport by substrates of the glucose transport system in *Escherichia coli*. Biochim. Biophys. Acta - Biomembr. 135: 1030–1051.

15. Aggarwal, R.K., and A. Narang. 2019. Deconstructing glucose-mediated catabolite repression of the *lac* operon of *Escherichia coli*: I. Inducer exclusion, by itself, cannot account for the repression. bioRxiv.: 739458.

16. Narang, A., A. Konopka, and D. Ramkrishna. 1997. The dynamics of microbial growth on mixtures of substrates in batch reactors. J. Theor. Biol. 184: 301–317.

17. Narang, A. 1998. The dynamical analogy between microbial growth on mixtures of substrates and population growth of competing species. Biotechnol. Bioeng. 59: 116–121.

18. Narang, A. 2006. Comparative analysis of some models of gene regulation in mixed-substrate microbial growth. J. Theor. Biol. 242: 489–501.

19. Narang, A., and S.S. Pilyugin. 2007. Bacterial gene regulation in diauxic and non-diauxic growth. J. Theor. Biol. 244: 326–48.

20. Cohn, M., and K. Horibata. 1959. Inhibition by glucose of the induced synthesis of the β-galactoside-enzyme system of *Escherichia coli*. Analysis of maintenance. J. Bacteriol. 78: 601–12.

21. Novick, A., and M. Weiner. 1957. Enzyme Induction as an All-or-None Phenomenon. Proc. Natl. Acad. Sci. 43: 553–566.

22. Ozbudak, E.M., M. Thattai, H.N. Lim, B.I. Shraiman, and A. Van Oudenaarden. 2004. Multistability in the lactose utilization network of *Escherichia coli*. Nature. 427: 737–40.

23. Savageau, M.A. 2001. Design principles for elementary gene circuits: Elements, methods, and examples. Chaos. 11: 142–159.

24. van Hoek, M.J.A., and P. Hogeweg. 2006. In silico evolved *lac* operons exhibit bistability for artificial inducers, but not for lactose. Biophys. J. 91: 2833–2843.

25. Dreisigmeyer, D.W., J. Stajic, M.E. Wall, I. Nemenman, and W.S. Hlavacek. 2008. Determinants of bistability in induction of the *Escherichia coli lac* operon. IET Syst. Biol. 2: 293–303.

26. Huber, R.E., J. Lytton, and E.B. Fung. 1980. Efflux of β-galactosidase products from *Escherichia coli*. J. Bacteriol. 141: 528–33.

27. Narang, A., and S.S. Pilyugin. 2008. Bistability of the *lac* operon during growth of *Escherichia coli* on lactose and lactose+glucose. Bull. Math. Biol. 70: 1032–64.

28. Savageau, M.A. 2011. Design of the *lac* gene circuit revisited. Math. Biosci. 231: 19–38.

29. Maharjan, R.P., and T. Ferenci. 2003. Global metabolite analysis: the influence of extraction methodology on metabolome profiles of *Escherichia coli*. Anal. Biochem. 313: 145–54.

30. Faijes, M., A.E. Mars, and E.J. Smid. 2007. Comparison of quenching and extraction methodologies for metabolome analysis of *Lactobacillus plantarum*. Microb. Cell Fact. 6: 27.

31. Bolten, C.J., P. Kiefer, F. Letisse, J.-C. Portais, and C. Wittmann. 2007. Sampling for metabolome analysis of microorganisms. Anal. Chem. 79: 3843–9.

32. Canelas, A.B., C. Ras, A. Pierick, J.C. Dam, J.J. Heijnen, and W.M. van Gulik. 2008. Leakage-free rapid quenching technique for yeast metabolomics. Metabolomics. 4: 226–239.

33. van Gulik, W.M. 2010. Fast sampling for quantitative microbial metabolomics. Curr. Opin. Biotechnol. 21: 27–34.

34. Taymaz-Nikerel, H., M. de Mey, C. Ras, A. ten Pierick, R.M. Seifar, J.C. van Dam, J.J. Heijnen, and W.M. van Gulik. 2009. Development and application of a differential method for reliable metabolome analysis in *Escherichia coli*. Anal. Biochem. 386: 9–19.

35. Tillack, J., N. Paczia, K. Nöh, W. Wiechert, and S. Noack. 2012. Error Propagation Analysis for Quantitative Intracellular Metabolomics. Metabolites. 2: 1012–1030.

36. Lancaster, J.R., R.J. Hill, and W.G. Struve. 1975. The characterization of energized and partially de-energized (respiration-independent) β-galactoside transport into *Escherichia coli*. Biochim. Biophys. Acta. 401: 285–98.

37. You, C., H. Okano, S. Hui, Z. Zhang, M. Kim, C.W. Gunderson, Y.-P. Wang, P. Lenz, D. Yan, and T. Hwa. 2013. Coordination of bacterial proteome with metabolism by cyclic AMP signalling. Nature. 500: 301–306.

38. Miller, J.H. 1972. Experiments in molecular genetics. Cold Spring Harbor Laboratory, Cold Spring Harbor, N.Y.

39. Cohn, M., and K. Horibata. 1959. Analysis of the differentiation and of the heterogeneity within a population of *Esherichia coli* undergoing induced β-galactosidase synthesis. J. Bacteriol. 78: 613–23.

40. Splechtna, B., T.-H. Nguyen, M. Steinböck, K.D. Kulbe, W. Lorenz, and D. Haltrich. 2006. Production of prebiotic galacto-oligosaccharides from lactose using β-galactosidases from *Lactobacillus reuteri*. J. Agric. Food Chem. 54: 4999–5006.

41. Cataldi, T.R.I., C. Campa, and G.E. De Benedetto. 2000. Carbohydrate analysis by high-performance anion-exchange chromatography with pulsed amperometric detection: The potential is still growing. Fresenius. J. Anal. Chem. 368: 739–758.

42. Bhattacharya, M., L. Fuhrman, A. Ingram, K.W. Nickerson, and T. Conway. 1995. Single-run separation and detection of multiple metabolic intermediates by anion-exchange high-performance liquid chromatography and application to cell pool extracts prepared from *Escherichia coli*. Anal. Biochem. 232: 98–106.

43. Koch, A.L. 2007. Growth Measurement. In: Methods for General and Molecular Microbiology, Third Edition. American Society of Microbiology. pp. 172–199.

44. Wright, J.K., I. Riede, and P. Overath. 1981. Lactose carrier protein of *Escherichia coli*: interaction with galactosides and protons. Biochemistry. 20: 6404–6415.

45. Kremling, A., J. Geiselmann, D. Ropers, and H. de Jong. 2015. Understanding carbon catabolite repression in *Escherichia coli* using quantitative models. Trends Microbiol. 23: 99–109.

46. Okano, H., R. Hermsen, K. Kochanowski, and T. Hwa. 2020. Regulation underlying hierarchical and simultaneous utilization of carbon substrates by flux sensors in *Escherichia coli*. Nat. Microbiol. 5: 206–215.

47. Huber, R.E., R. Pisko-Dubienski, and K.L. Hurlburt. 1980. Immediate stoichiometric appearance of β-galactosidase products in the medium of *Escherichia coli* cells incubated with lactose. Biochem. Biophys. Res. Commun. 96: 656–661.

48. Winkler, H.H., and T.H. Wilson. 1966. The role of energy coupling in the transport of β-galactosides by *Escherichia coli*. J. Biol. Chem. 241: 2200–2211.

49. Egel, R. 1988. The “*lac*” operon: an irrelevant paradox? Trends Genet. 4: 31.

50. Marbach, A., and K. Bettenbrock. 2012. *lac* operon induction in *Escherichia coli*: Systematic comparison of IPTG and TMG induction and influence of the transacetylase LacA. J. Biotechnol. 157: 82–8.

51. Herzenberg, L.A. 1959. Studies on the induction of β-galactosidase in a cryptic strain of *Escherichia coli*. Biochim. Biophys. Acta. 31: 525–538.

52. Milo, R., and R. Phillips. 2015. Chapter 4: Rates and Durations. In: Cell biology by the numbers. Garland Science, New York. pp. 209–282.

53. Huber, R.E., K. Wallenfels, and G. Kurz. 1975. The action of β-galactosidase (*Escherichia coli*) on allolactose. Can. J. Biochem. 53: 1035–8.

54. Huber, R.E., and K.L. Hurlburt. 1984. *Escherichia coli* growth on lactose requires cycling of β-galactosidase products into the medium. Can. J. Microbiol. 30: 411–5.

55. Kuhlman, T., Z. Zhang, M.H. Saier, and T. Hwa. 2007. Combinatorial transcriptional control of the lactose operon of *Escherichia coli*. Proc. Natl. Acad. Sci. U. S. A. 104: 6043–8.

56. Huber, R.E., G. Kurz, and K. Wallenfels. 1976. A quantitation of the factors which affect the hydrolase and transgalactosylase activities of β-galactosidase (*E. coli*) on lactose. Biochemistry. 15: 1994–2001.

