## Supplemental for "Deconstructing glucose-mediated catabolite repression of the *lac* operon of *Escherichia coli*: II. Positive feedback exists and drives the repression"

**Supplement**

### S1 Mathematical theory motivating this work

We have hypothesized that the *lac* operon is subject to positive feedback because (a) the inducer stimulates the synthesis of the lactose enzymes, and (b) the lactose enzymes in turn promote the accumulation of the inducer. However, based on mathematical models (23–25), it has been argued that the foregoing positive feedback loop exists in the presence of TMG but not lactose, and this is because both TMG and allolactose stimulate synthesis of the lactose enzymes, but the lactose enzymes promote the accumulation of only TMG, and not allolactose.

The first goal of this section is to show that the above conclusion is based upon the following three assumptions:

1. Intracellular TMG is not metabolized — it is excreted from the cell by diffusion.
2. Intracellular lactose and allolactose are not excreted — they are metabolized by the β-galactosidase (LacZ).
3. The permease (LacY) activity is proportional to the LacZ activity.

To this end, we revisit the models to show that under these assumptions, positive feedback exists in the presence of TMG, but not lactose. Now, as discussed in the Introduction, assumptions 2 and 3 are not supported by the data. Thus, the second goal of this section is to show that relaxing either one of these two assumptions restores positive feedback in the presence of lactose.

#### S1.1 Induction of *lac* operon in the presence of TMG

In this case, it is assumed that (Fig. S1)

1. TMG is imported into the cell by the permease at the rate

$$r_{T,p}^{+} = k_{T,p}^{+} E_p \frac{T_e}{K_{T,p}^{+} + T_e} = k_{T,p}^{+} \left(\frac{E_p}{E_g}\right) E_g \frac{T_e}{K_{T,p}^{+} + T_e}, \qquad \text{S1}$$

where $E_p$ and $E_g$ denote the specific permease and β-galactosidase activities, and $T_e$ denotes the extracellular TMG concentration. We have replaced $E_p$ by $(E_p/E_g)E_g$ because this will facilitate the subsequent consideration of two special cases, namely, constant and increasing $E_p/E_g$.

2. TMG is exported from the cell by passive diffusion, i.e.,

$$r_{T,d}^{-} = k_{T,d}^{-}(T - T_e), \qquad \text{S2}$$

where $T$ denotes the intracellular TMG concentration.

Under these assumptions, the evolution of $T$ is governed by the equation

$$\frac{dT}{dt} = r_{T,p}^{+} - r_{T,d}^{-} - \mu T = k_{T,p}^{+} \left(\frac{E_p}{E_g}\right) E_g \frac{T_e}{K_{T,p}^{+} + T_e} - k_{T,d}^{-}(T - T_e) - \mu T, \qquad \text{S3}$$

where $\mu T$ denotes the dilution of intracellular TMG due to expansion of the cells by growth. Now, intracellular TMG accumulates and depletes on a time scale of minutes (Fig. S3 of (15)), whereas lactose enzymes are synthesized and diluted on a time scale of hours. It follows that during the slow evolution of the lactose enzymes, the intracellular TMG concentrations are in a quasi-steady state which satisfies the equation

$$0 \approx \frac{dT}{dt} \approx r_{T,p}^{+} - r_{T,d}^{-} \Longrightarrow T \approx T_e + \frac{k_{T,p}^{+}}{k_{T,d}^{-}} \left(\frac{E_p}{E_g}\right) E_g \frac{T_e}{K_{T,p}^{+} + T_e}, \qquad \text{S4}$$

where TMG dilution is neglected since it is small compared to TMG expulsion. If $E_p$ is proportional to $E_g$, then Eq. (S4) implies that the quasi-steady state concentration of intracellular TMG increases linearly with the induction level $E_g$. Alternatively, one can say that the lactose enzymes promote the accumulation of intracellular TMG.

### S1.2 Induction of *lac* operon in the presence of lactose

We shall now consider if lactose enzymes promote the accumulation of intracellular allolactose during growth on lactose. In this case, it is assumed that (Fig. S2)

1. Lactose is imported into the cell by the permease at the rate

$$r_{L,p}^{+} = k_{L,p}^{+} E_p \frac{L_e}{K_{L,p}^{+} + L_e} = k_{L,p}^{+} \left(\frac{E_p}{E_g}\right) E_g \frac{L_e}{K_{L,p}^{+} + L_e}, \tag{S5}$$

   where $L_e$ denotes the extracellular lactose concentration.

2. Intracellular lactose is exported from the cell by passive diffusion, i.e.,

$$r_{L,d}^{-} = k_{L,d}^{-}(L - L_e), \tag{S6}$$

   where $L$ denotes the intracellular lactose concentration.

However, since intracellular lactose is metabolizable, it also undergoes two additional reactions catalysed by β-galactosidase:

3. Intracellular lactose is hydrolysed to produce glucose and galactose at the rate

$$r_{L,h} = k_{L,h} E_g \frac{L}{K_{L,h} + L}, \tag{S7}$$

   where $E_g$ denotes the specific β-galactosidase activity.

4. Intracellular lactose is transgalactosylated to produce the inducer, allolactose, at the rate

$$r_{L,t} = k_{L,t} E_g \frac{L}{K_{L,t} + L}. \tag{S8}$$

The intracellular allolactose thus produced has two possible fates:

5. Intracellular allolactose is expelled from the cell by passive diffusion at the rate

$$r_{A,d}^{-} = k_{A,d}^{-}(A - A_e), \tag{S9}$$

where $A$ and $A_e$ denote the intracellular and extracellular allolactose concentrations, respectively.

6. Intracellular allolactose is also hydrolysed by β-galactosidase to produce glucose and galactose at the rate

$$r_{A,h} = k_{A,h} E_g \frac{A}{K_{A,h} + A}, \tag{S10}$$

It follows that the evolution of intracellular lactose and allolactose is governed by the equations

$$\frac{dL}{dt} = r_{L,p}^{+} - r_{L,d}^{-} - r_{L,h} - r_{L,t} - \mu L, \tag{S11}$$

$$\frac{dA}{dt} = r_{L,t} - r_{A,d}^{-} - r_{A,h} - \mu A. \tag{S12}$$

Again, we are concerned with phenomena, such as induction and repression of lactose enzymes, which occur on the time scale of enzyme synthesis and dilution (1 h), but metabolites, such as intracellular lactose and allolactose, turn over on a time scale of 1 s (52). It follows that during the slow evolution of the lactose enzymes in induction and repression, the concentrations of intracellular lactose and allolactose are in a quasi-steady state satisfying the equations

$$0 \approx \frac{dL}{dt} \approx r_{L,p}^{+} - r_{L,d}^{-} - r_{L,h} - r_{L,t}, \tag{S13}$$

$$0 \approx \frac{dA}{dt} \approx r_{L,t} - r_{A,d}^{-} - r_{A,h}. \tag{S14}$$

In what follows, we shall use these equations to determine, under various limiting conditions, the variation of the quasi-steady state concentration of intracellular allolactose with the induction level.

#### S1.2.1 There is no positive feedback if metabolism dominates expulsion

We begin by considering the limiting case wherein metabolism dominates expulsion (i.e., $r_{L,d}^{-} \ll r_{L,h}, r_{L,t}$ and $r_{A,d}^{-} \ll r_{A,h}$), which was assumed in the mathematical models showing the absence of positive feedback in the presence of lactose (23–25). Under these conditions, Eqs. (S13)–(S14) have the form

$$0 \approx \frac{dL}{dt} \approx r_{L,p}^{+} - r_{L,h} - r_{L,t}$$
$$= k_{L,p}^{+}\left(\frac{E_p}{E_g}\right)E_g\frac{L_e}{K_{L,p}^{+} + L_e} - k_{L,h}E_g\frac{L}{K_{L,h} + L} - k_{L,t}E_g\frac{L}{K_{L,t} + L}, \quad \text{S15}$$

$$0 \approx \frac{dA}{dt} \approx r_{L,t} - r_{A,h} = k_{L,t}E_g\frac{L}{K_{L,t} + L} - k_{A,h}E_g\frac{A}{K_{A,h} + A}. \quad \text{S16}$$

Now if $E_p$ is proportional to $E_g$, Eq. (S15) implies that the quasi-steady state concentration of intracellular lactose $L$ is independent of the induction level $E_g$ — it is completely determined by the extracellular lactose concentration $L_e$, i.e., $L = L(L_e)$. Furthermore, Eq. (S16) implies that the quasi-steady state concentration of intracellular allolactose $A$ is also independent of the induction level $E_g$ — it is completely determined by the quasi-steady state intracellular lactose concentration, i.e., $A = A(L)$. It follows that $A = A(L(L_e))$ is independent of the induction level $E_g$, i.e., the lactose enzymes fail to promote *any* accumulation of intracellular allolactose.

#### S1.2.2 Positive feedback can exist if there is significant expulsion of lactose/allolactose

The foregoing analysis neglected the existence of lactose and allolactose expulsion from the cells (26, 47, 53, 54). To understand the effect of expulsion, it is useful to examine the limiting case wherein expulsion dominates metabolism (i.e., $r_{L,d}^{-} \gg r_{L,h}, r_{L,t}$ and $r_{A,d}^{-} \gg r_{A,h}$). In this case, Eqs. (S13)–(S14) have the form

$$0 \approx \frac{dL}{dt} \approx r_{L,p}^{+} - r_{L,d}^{-} = k_{L,p}^{+}\left(\frac{E_p}{E_g}\right)E_g\frac{L_e}{K_{L,p}^{+} + L_e} - k_{L,d}^{-}(L - L_e), \quad \text{S17}$$

$$0 \approx \frac{dA}{dt} \approx r_{L,t} - r_{A,d}^{-} = r_{L,t} = k_{L,t} E_g \frac{L}{K_{L,t} + L} - k_{A,d}^{-}(A - A_e), \tag{S18}$$

which imply that

$$L \approx L_e + \frac{k_{L,p}^{+}}{k_{L,d}^{-}} \left(\frac{E_p}{E_g}\right) E_g \frac{L_e}{K_{L,p}^{+} + L_e}, \tag{S19}$$

$$A = A_e + \frac{k_{L,t}}{k_{A,d}^{-}} E_g \frac{L}{K_{L,t} + L}. \tag{S20}$$

Now Eq. (S19) implies that if $E_p$ is proportional to $E_g$, then $L$ increases linearly with the induction level $E_g$. This equation is formally identical to Eq. (S4) because under the assumed condition (expulsion dominates metabolism), lactose and TMG suffer identical fates. Substituting (S19) in (S20) implies that $A$ increases with $E_g$ faster than linearly. This sigmoidal variation of $A$ reflects the combined effect of two steps. Indeed, if expulsion dominates metabolism of *both* $L$ and $A$, then both steps contribute to ensure that $A$ increases with $E_g$ faster than linearly. It is clear, however, that $A$ increases linearly with $E_g$ if expulsion dominates metabolism of *either* $L$ *or* $A$, and hence, positive feedback can exist if either lactose or allolactose undergoes significant expulsion.

We have shown above that there is no positive feedback when metabolism dominates expulsion, and strong positive feedback when expulsion dominates metabolism. Experiments show that both lactose and allolactose are expelled (26, 47). Since the relative rates of lactose and allolactose metabolism and expulsion are not known, we cannot rule out the possibility that lactose or/and allolactose expulsion are large enough to cause significant positive feedback.

#### S1.2.3 Positive feedback can exist if there is inducer exclusion

Thus far, we have assumed that $E_p$ is proportional to $E_g$, but this assumption is not true in the presence of inducer exclusion (15). To see this, observe that due to coordinate synthesis of the permease and β-galactosidase, the number of β-galactosidase molecules synthesized is proportional to the *total* number of permease molecules synthesized, but a certain fraction is inactivated in the presence of inducer exclusion. Since this fraction varies with the induction level, so does the LacY:LacZ ratio. More precisely, if we denote the concentration of *all* permease molecules by $E_{p,t}$, and the concentration of *active* permease molecules by $E_p$, then

$$\frac{E_p}{E_g} = \frac{E_p}{E_{p,t}}\frac{E_{p,t}}{E_g}. \qquad \text{S21}$$

Now, $E_{p,t}$ is proportional to $E_g$ because of coordinate synthesis, but $E_p$ is proportional to $E_g$ precisely when $E_p/E_{p,t}$ is constant. This is certainly true in the absence of inducer exclusion since all the permease molecules are active ($E_p/E_{p,t} = 1$). However, in the presence of inducer exclusion, $E_p/E_{p,t}$ increases with $E_g$ since the more induced the cells, the larger the fraction of active permease molecules $E_p/E_{p,t}$ (Fig. 2 of (15)). It follows that in the presence of inducer exclusion, $E_p/E_g$ is not constant, but increases with $E_g$.

If $E_p/E_g$ increases with $E_g$, as is the case in the presence of inducer exclusion, then $A$ is an increasing function of the induction level $E_g$ even if metabolism dominates expulsion. Indeed, under this condition, Eq. (S15) implies that the quasi-steady concentrations of intracellular lactose, $L = L(E_p/E_g, L_e)$, increases with the induction level $E_g$. It follows from Eq. (S16) that the quasi-steady state concentration of allolactose, $A = A(L)$, also increases with the induction level $E_g$, thus implying the potential for positive feedback in the presence of inducer exclusion.

**S2 Consumption of extracellular allolactose is mediated by Lac permease**

To determine if allolactose uptake was mediated by LacY, we studied the differential rate of allolactose uptake in a *lacY*⁻ strain. Before discussing this data, it is useful to recall that the rates of allolactose expulsion and uptake satisfy the relation (see Eq. 6)

$$r_{A,d}^{-} - r_A^{+} = \mu \frac{dA_e}{dX}, \quad \text{S22}$$

where $\mu$ is the specific growth rate, $r_{A,d}^{-}$ is the specific allolactose efflux rate, $r_A^{+}$ is the specific allolactose uptake rate, and $dA_e/dX$ is the slope of the differential plot. We have shown above that when pre-induced wild-type cells are exposed to a mixture of lactose (4 mM) and maltose (2.8 mM), $dA_e/dX$ decreases monotonically because $r_{A,d}^{-}$ is constant and $r_A^{+}$ grows progressively due to the accumulation of allolactose in the medium. We are now concerned with the outcome of the same experiment performed with pre-induced *lacY*⁻ cells.

Now, if uptake of allolactose and lactose uptake were mediated solely by the permease, there would no uptake of lactose and allolactose in *lacY*⁻ cells. Consequently, there would be neither allolactose efflux ($r_{A,d}^{-} = 0$) nor allolactose uptake ($r_A^{+} = 0$), and hence, the slope of the differential plot $dA_e/dX$ would be zero. However, when pre-induced *lacY*⁻ cells were exposed to a mixture of lactose (4 mM) and maltose (2.8 mM), the differential plot had a positive, but constant, slope (○ in Fig. S4).

The positive slope implies that lactose uptake was not mediated solely by the permease — it entered the cell by diffusion, where it was hydrolysed to produce intracellular allolactose, part of which accumulates in the medium. We confirmed that lactose uptake in *lacY*⁻ cells was mediated primarily by diffusion because when the above experiment was performed with a 10-fold smaller concentration of lactose (0.4 mM lactose + 2.8 mM maltose), the slope of the differential plot decreased 7-fold (△ in Fig. S4).

Although allolactose is produced and expelled by *lacY*⁻ cells, its consumption is negligibly small compared to that observed in wild-type cells. Indeed, given the constant specific growth rate and the slope of the differential plot for the data labelled ○ in Fig. S4, we calculated that $r_{A,d}^{-} - r_A^{+} = 15$ μmols gdw$^{-1}$ h$^{-1}$. Since allolactose accumulates in the medium, $r_A^{+}$ must be increasing, but since $r_{A,d}^{-} - r_A^{+}$ remains constant, the magnitude of $r_A^{+}$ must always remain negligible compared to the constant slope of 15 μmols gdw$^{-1}$ h$^{-1}$. It follows that $r_A^{+} \lesssim 1.5$ μmols gdw$^{-1}$ h$^{-1}$ even though the extracellular concentrations of allolactose became as high as 5 μM. At this concentration, the allolactose uptake rate in wild-type cells is 40 μmols gdw$^{-1}$ h$^{-1}$ (Fig. 3B). We conclude that $r_A^{+}$ is reduced at least 25-fold in $lacY^{-}$ cells.

**S3 The normalized induction curves for allolactose and IPTG coincide.**

Since *lac* expression is modulated by cAMP and inducer, the induction rate can be written as $r_{E_g} = r_{E_g}(C, I)$, where denoted $C$ and $I$ denote the concentrations of cAMP and inducer. If the effects of cAMP and inducer are independent, the induction rate has the form $r_{E_g} = V_{E_g} f_C(C) f_I(I)$, where $V_{E_g}$ denotes the maximum induction rate, and $0 < f_C, f_I < 1$ are increasing functions approaching 1 at sufficiently high cAMP and inducer concentrations. It follows that if the induction rate $r_{E_g}$ is normalized by the induction rate $V_{E_g} f_C(C)$ obtained under fully induced conditions ($f_I(I) \to 1$), this normalized induction rate is completely determined by the inducer concentration, i.e,

$$\rho_{E_g} \triangleq \frac{r_{E_g}}{V_{E_g} f_C(C)} = f_I(I). \qquad \text{S23}$$

Now, mechanistic models of *lac* induction show that $\rho_{E_g}$ always has the functional form

$$\rho_{E_g} = f_I(I) = f\left(\frac{I}{K_I}\right), \qquad \text{S24}$$

where $K_I$ denotes the dissociation constant for binding of the inducer to the Lac repressor, i.e., the normalized induction curves for all inducers are the same except for differences in the scales corresponding to their abscissae (19, 55). In particular, the normalized induction rates obtained in the presence of allolactose and IPTG have the form

$$\rho_{E_g} = f\left(\frac{A}{K_A}\right), \rho_{E_g} = f\left(\frac{[\mathrm{IPTG}]}{K_{\mathrm{IPTG}}}\right). \qquad \text{S25}$$

Since $K_A = K_{\mathrm{IPTG}} = 0.1$ μM (2), the normalized induction curves for allolactose and IPTG coincide.

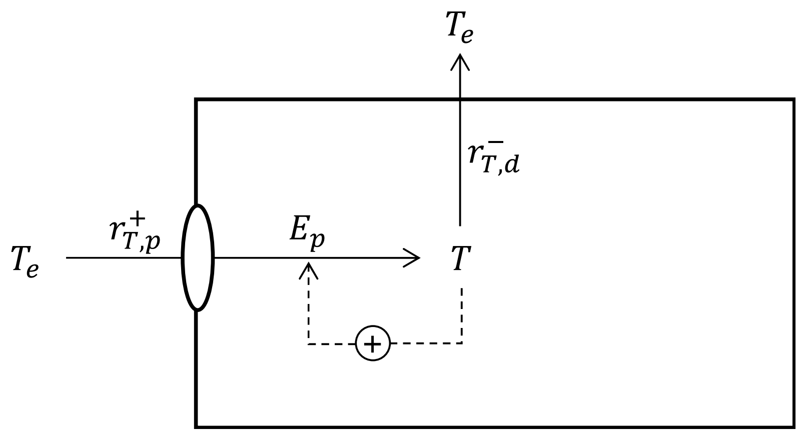


Fig. S1: Fate of TMG in *E. coli*: Extracellular TMG ($T_e$) is imported into the cell by permease ($E_p$), and intracellular TMG ($T$) is exported out of the cell by diffusion. The dashed line represents induction of permease by intracellular TMG.

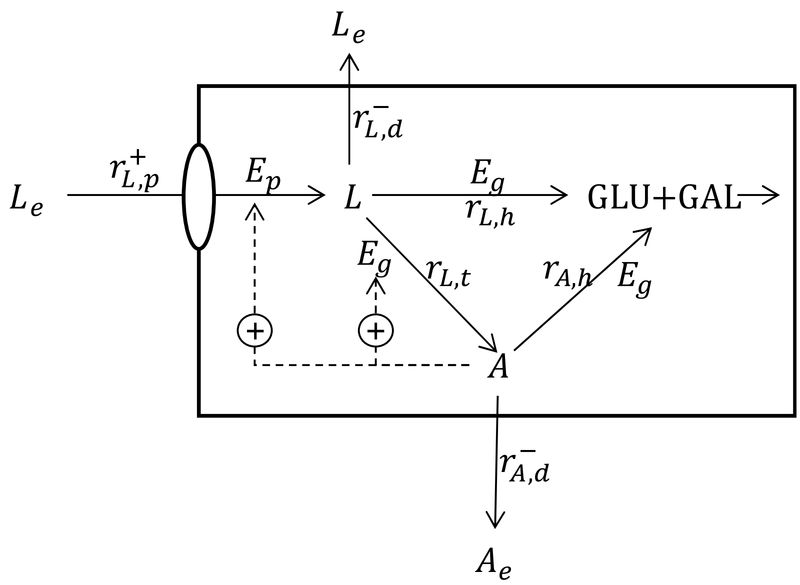


Fig. S2: Fate of lactose in *E. coli* (adapted from (56)): Extracellular lactose ($L_e$) is imported into the cell by permease ($E_p$). The intracellular lactose ($L$) thus produced is exported from the cell by passive diffusion, and converted by β-galactosidase ($E_g$) to glucose + galactose and allolactose ($A$). Intracellular allolactose is hydrolysed to glucose + galactose by β-galactosidase, and exported from the cell by passive diffusion thus producing extracellular allolactose ($A_e$). The dashed lines represent induction of permease and β-galactosidase by allolactose.

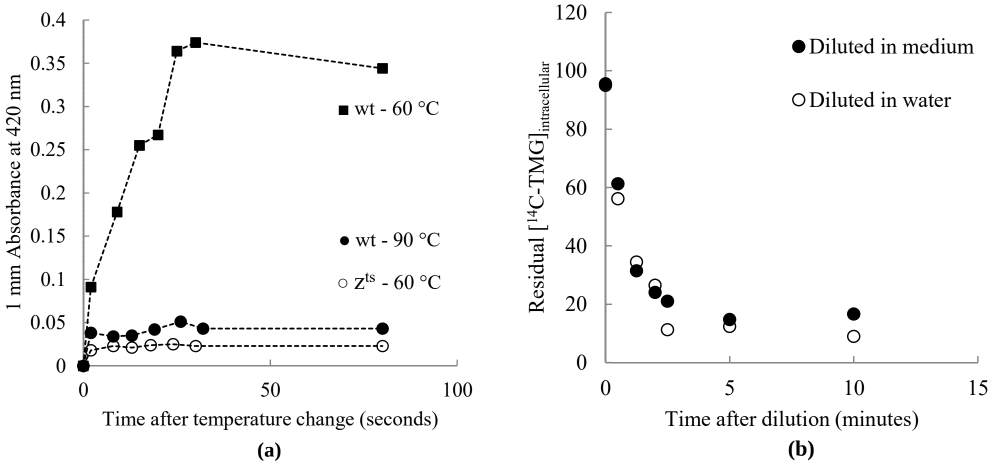


Fig. S3: Heating rapidly deactivates β-galactosidase, but significant efflux occurs. (A) One volume of wild-type (wt) or mutant cells with temperature-sensitive LacZ ($z^{ts}$)), pre-induced with 0.5 mM IPTG at 30 °C, were permeabilized and added at t=0 to five volumes of rapidly mixed ONPG solution pre-heated to the indicated temperature. Absorbance of the resulting colour change was measured at 420 nm. (B) Wild-type cells pre-loaded with [$^{14}$C]TMG were diluted at t=0 in pre-warmed fresh medium or water devoid of TMG. Equal volumes of cells were filtered at various times and disintegrations per unit time were measured and reported as intracellular TMG concentration.

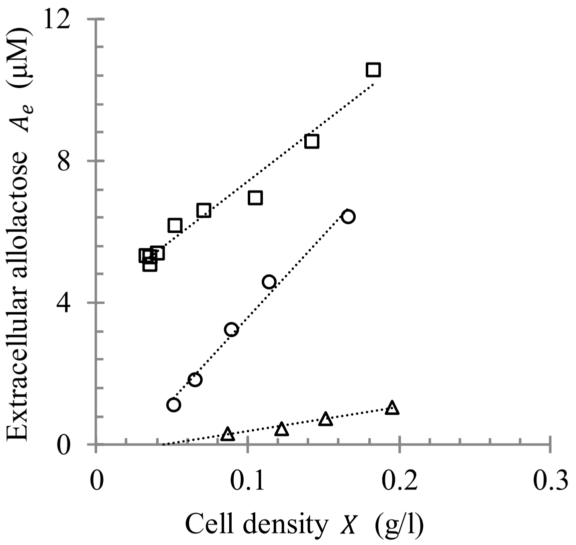


Fig. S4: Differential plots showing the variation of the extracellular allolactose concentration as a function of the biomass concentration of the permease-deficient strain RA-Y66. Cells were pre-grown on maltose and 0.1 mM IPTG to achieve similar hydrolytic activity of β-galactosidase. To start the experiment, permease-deficient cells were re-suspended in medium containing 2.8 mM maltose and varying combinations of lactose and allolactose concentrations: 4 mM lactose (○), 0.4 mM lactose (▵), and 4 mM lactose + 5 μM allolactose (□). The cell density, measured as optical density at 600 nm, was to calculate the specific growth rate μ in each case, and the extracellular allolactose concentration was measured in the filtrate using ion-exchange chromatography. The dotted lines show the linear fits used to compute the slopes.

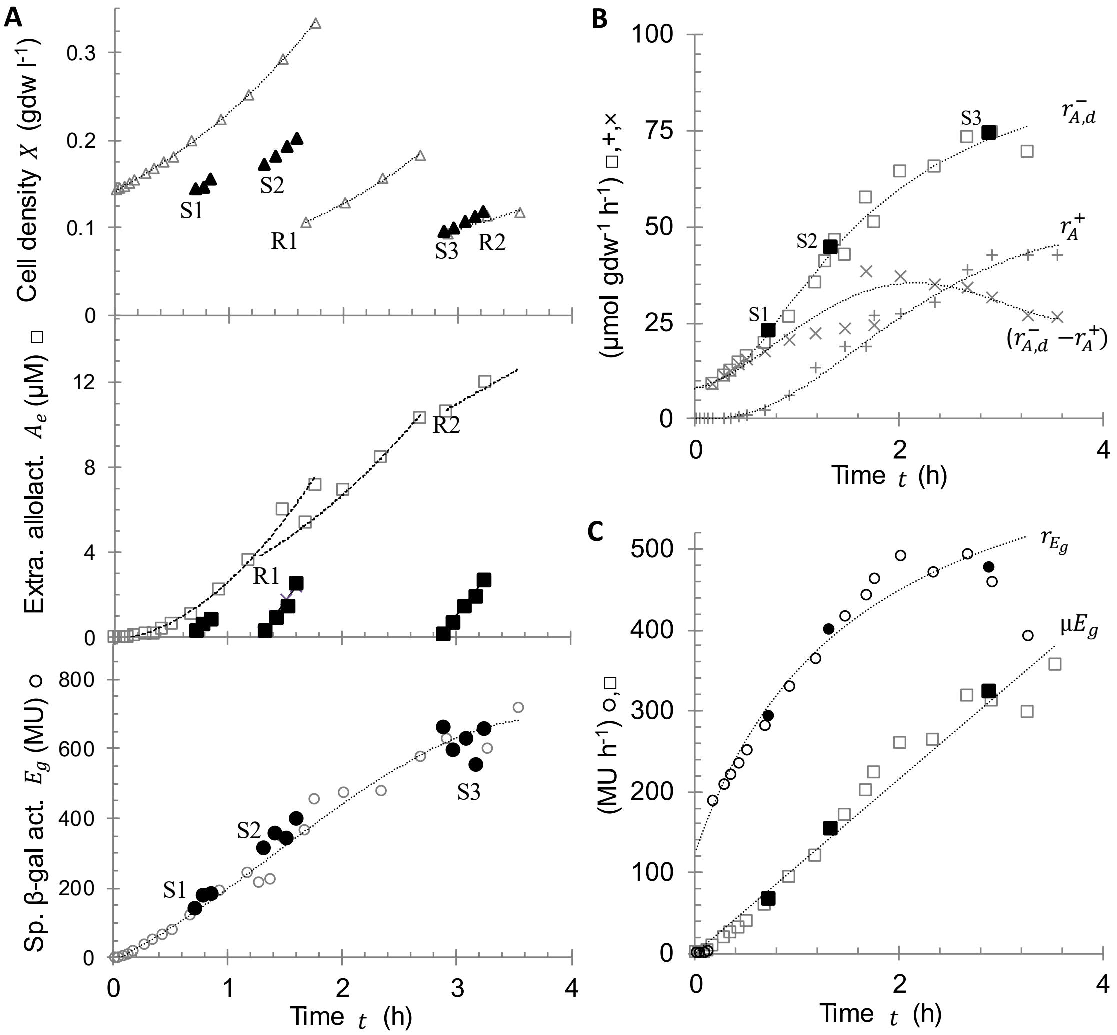


Fig. S5: Determination of the instantaneous specific allolactose efflux rate $r_{A,d}^-(t)$, induction rate $r_{E_g}(t)$, and dilution rate $\mu(t)E_g(t)$ during growth of an initially uninduced culture of *E. coli* K12 MG1655 on lactose + maltose. (A) Evolution of the cell density $X$ (△), extracellular allolactose concentration $A_e$ (□), and specific β-galactosidase activity $E_g$ (○). To keep the cell density sufficiently low, the culture was centrifuged at the points labelled R1, R2, and a fraction of the cells were reintroduced into the same medium. To determine the initial allolactose uptake rates, the cells were transferred to fresh medium at the time points labelled S1, S2, S3, after which $X$ (▲), $A_e$ (■), and $E_g$ (●) and were measured for 10–20 min. (B) Evolution of the semi-empirical (□) and empirical (■) $r_{A,d}^-(t)$ derived from the data in (A). (C) Evolution of $r_{E_g}(t)$ and $\mu(t)E_g(t)$ derived from the data in (A).

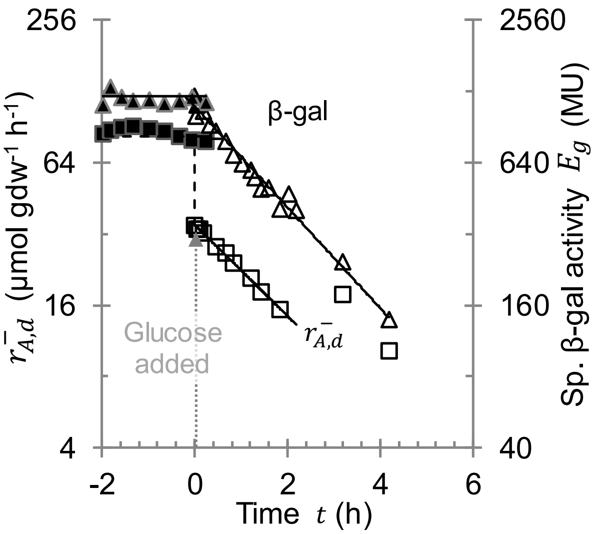


Fig. S6: Repression kinetics of fully induced *E. coli* K12 MG1655 cells upon transfer to glucose + lactose. At $t = 0$, cells growing exponentially on maltose (2.8 mM) + lactose (4 mM) were transferred to a pre-warmed medium containing glucose (2.1 mM) + lactose (4 mM), and the subsequent specific allolactose efflux rate $r_{A,d}^{-}$ (Δ) and specific β-glactosidase activity $E_g$ (□) were measured. Data for four generation times are shown.

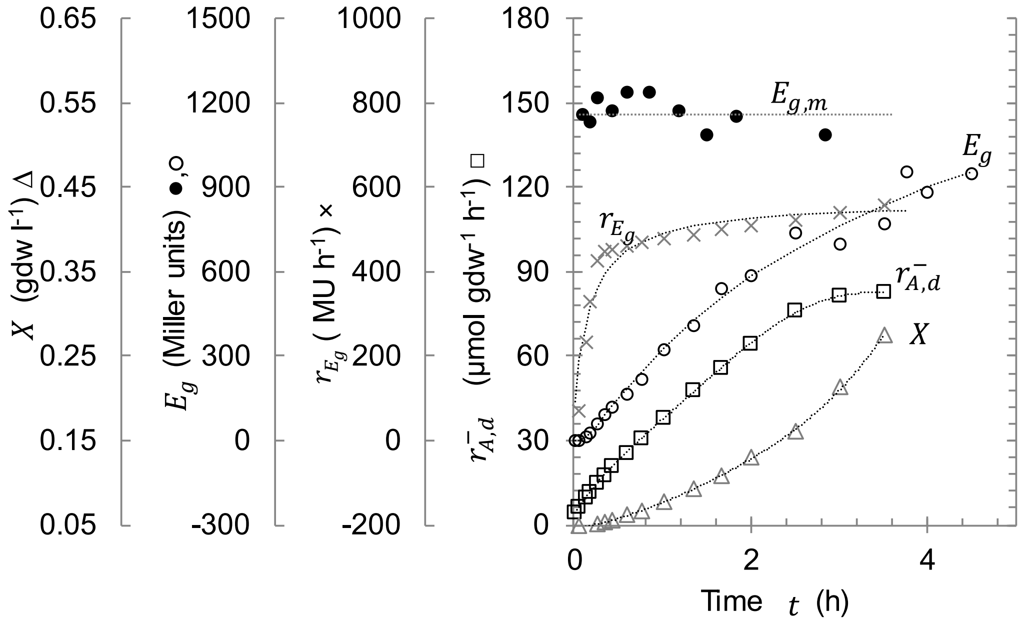


Fig. S7: The evolution of the induction rate $r_{E_g}$ and the specific allolactose efflux rate $r_{A,d}^{-}$during growth of initially uninduced *E. coli* K12 MG1655 cells on lactose + maltose.

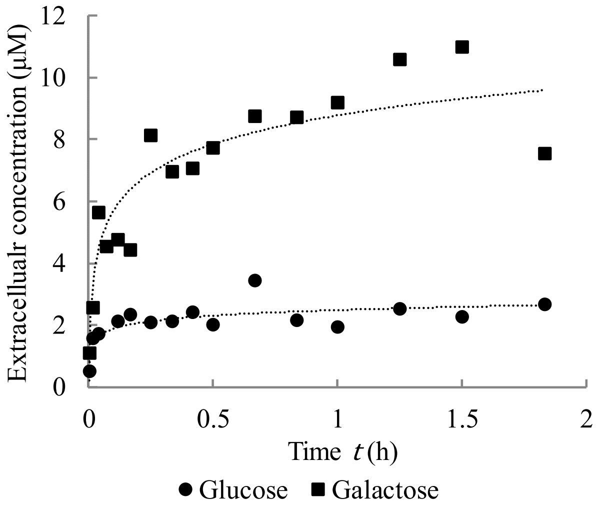


Fig. S8: Extracellular accumulation of galactose and glucose in the medium. Cells pre-grown on lactose and maltose were spun and resuspended in fresh medium containing 4 mM lactose and 2.8 mM maltose. The extracellular concentrations of glucose and galactose were measured chromatographically, and used to calculate the initial efflux rates of galactose and glucose reported in Table 1.

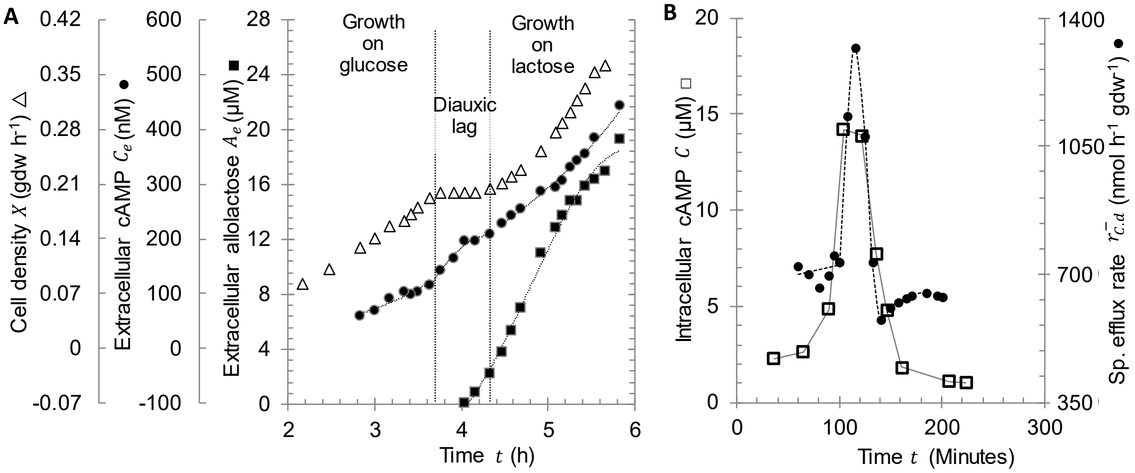


Fig – No bookmark: The specific cAMP efflux rate is proportional to the intracellular cAMP concentration. (A) Evolution of the cell density and the concentrations of extracellular cAMP and allolactose. Cells pre-grown on 7.8 mM glucose were transferred to medium containing glucose 2.2 mM glucose and 4 mM lactose. (B) Comparison of specific cAMP efflux rates obtained from the data in (A) with published data for intracellular cAMP (□). The specific cAMP efflux rates, computed using equation (7) as described in the text, are proportional to the intracellular cAMP levels reported by Inada et al. (1996). In particular, the specific cAMP efflux rates before and after the diauxic lag are comparable, but increase transiently during the diauxic lag. The x-axis has been translated by 110 minutes for comparison.
